# Distinct sensorimotor encoding in tuft dendrites and somata associated with action, correction, and learning

**DOI:** 10.64898/2026.05.06.722323

**Authors:** Jackson Scheib, Zachary L Newman, Jacob Gable, Deano M Farinella, Mitchell Head, Savannah Bliese, Benjamin Dougen, Harishankar Jayakumar, Sarah Young, Nicole Miller, Robert Al Khoury, Huan Tran, Tien Dinh, Aaron Kerlin

## Abstract

Frontal cortex plays critical roles in action control and motor skill learning. Within the layer 1 apical tuft dendrites of layer 5 (L5) neurons in frontal cortex, precise input patterns and back-propagating action potentials can trigger powerful regenerative events that may be essential for flexible computation and learning. However, it remains unclear whether tuft activity in frontal cortical L5 circuits encodes sensorimotor information that differs from the information conveyed by their outputs to downstream targets. Using longitudinal two-photon calcium imaging, we investigated sensorimotor encoding in the apical tuft dendrites and somata of L5 extratelencephalic (ET) neurons in the frontal cortex of mice during learning of a discrete change to a cued dexterous action. During learning, movement errors either triggered corrective action or did not, allowing us to dissociate error signals from signals selective for corrective action. Somatic activity tracked both instructional cues and action, whereas tuft activity predominantly tracked instructional cues. Movement errors during learning revealed additional distinct tuft activity that was selectively associated with corrective actions. Furthermore, learning induced divergent changes in the response gain and net selectivity of tuft dendrites compared to somata. Our measurements uncover systematic differences between the tuft dendrites and somata in sensorimotor selectivity, sensitivity to corrective action, and functional plasticity, providing a foundation for investigating the contributions of dendritic computation to motor skill learning.

## Introduction

During the acquisition and refinement of complex behavior, predictions about the state of the external world and the efficacy of actions are tested and revised. Take answering a phone as an example. From sensory input (*i.e.*, the ring tone) a latent aspect of the world (*i.e.*, ‘call being received’) is inferred, as is an optimal course of action (*i.e.*, ‘swipe upward until phone unlocks’). However, when an action unfolds, errors in the inferred state (e.g., it was actually your coworker’s phone) or inferred action (*e.g.*, a larger swipe is required to unlock) are identified, triggering corrective action and storage of information about the experience that will guide future estimates (*i.e.*, learning). The frontal cortex plays a critical role in such flexible motor behaviors (***Lopes et al., 2023***; ***Mizes et al., 2024***; ***Dragoi et al., 2026***; ***Dasgupta et al., 2025***) and corrective action (***Heindorf et al., 2018***; ***Bollu et al., 2021***), especially in the early stages of learning a new motor skill (***Kawai et al., 2015***; ***Hwang et al., 2019***).

The circuit mechanisms of flexible behavior and rapid learning may depend on the nonlinear integration of signals in the apical tuft dendrites of layer 5 (L5) cortical pyramidal neurons. The tuft dendrites of L5 neurons receive distinct synaptic inputs (***Spratling, 2002***; ***Petreanu et al., 2009***) and support regenerative events – tuft spikes – that depend on precise patterns of synaptic input and back-propagating action potentials (***Larkum et al., 1999a***; ***Larkum et al., 1999b***; ***Stuart and Häusser, 2001***; ***Larkum et al., 2009***; ***Farinella et al., 2014***; ***Stuart and Spruston, 2015***). These tuft spikes facilitate burst firing (***Larkum et al., 1999b***) and generate a large influx of calcium, impacting both the ongoing output and the future function of neurons. In sensory cortices, evidence indicates that tuft spikes condition the sensory responses of L5 somata on top-down contextual signals (***Larkum, 2013***; ***Xu et al., 2012***; ***Ranganathan et al., 2018***; ***Fişek et al., 2023***; ***Takahashi et al., 2016***; ***Zhang et al., 2025***) and can rapidly induce persistent changes in the selectivity of L5 neurons (***Xiao et al., 2025***; ***Yaeger et al., 2025***). Furthermore, during learning, activity within the tuft of L5 neurons in primary sensory cortex changes along sensory dimensions relevant to a newly learned task (***Benezra et al., 2024***).

Tuft activity may serve different functions in the frontal cortex, however. Intracortical and thalamic connectivity motifs involving the tuft are different in the frontal cortex compared to primary sensory cortices (***Shepherd and Yamawaki, 2021***; ***Geng et al., 2022***). These connectivity motifs may support long-timescale and rotational frontal cortex dynamics (***Murray et al., 2014***; ***Shepherd and Yamawaki, 2021***) related to action planning and online movement correction (***Riehle and Requin, 1989***; ***Churchland et al., 2006***; ***Kaufman et al., 2016***; ***Heindorf et al., 2018***; ***Ames et al., 2019***; ***Sauerbrei et al., 2020***). Tuft connectivity also changes during motor skill learning (***Xu et al., 2009***), concurrent with changes in the activity patterns of L5 neurons (***Costa et al., 2004***; ***Peters et al., 2017***). Recent models propose that tuft spikes may encode both action initiation information (***Takahashi et al., 2021***; ***Guzulaitis and Palmer, 2023***) and teaching information (***Rao, 2024***; ***Senn et al., 2024***) that are distinct from the final output of neurons encoded in somatic spiking. Changes in laminar dynamics throughout learning also differ between the frontal and sensory cortex (***Pollak et al., 2025***), suggesting that representations within L5 neurons could change in both a compartment-specific and area-specific manner. However, functional encoding in the tuft dendrites and somata has not been directly compared during motor learning. As a result, it remains unclear what distinct information, if any, is conveyed by spikes in the tuft dendrites of L5 neurons within the frontal cortex.

We addressed this knowledge gap by performing longitudinal two-photon (2P) calcium imaging of apical tuft dendrites and somata of L5 extratelencephalic (ET) neurons in anterolateral motor cortex (ALM), a premotor region in the frontal cortex of mice. L5 ET neurons play a central role in encoding the trajectory of upcoming and ongoing movement (***Evarts, 1968***; ***Li et al., 2015***) and are likely to be involved in the implementation of new dexterous actions (***Peters et al., 2017***; ***Li et al., 2017***; ***Serradj et al., 2023***). To measure activity associated with the acquisition of new motor trajectories, we trained mice to expert performance on a cued directional licking task and then shifted the lick targets. Relative to somatic activity, tuft activity showed weaker associations with the timing of the initial action but stronger associations with instructional cues and corrective actions, revealing compartmental differences that had not previously been reported. The gain of task-modulated activity within the two compartments also changed in opposite directions during learning. Our delineation of functional differences between tuft activity and somatic output provides critical new evidence to guide circuit models of credit assignment, motor control, and motor skill learning.

## Results

### Contingency-dependent motor skill learning

To better understand the role of tuft dendrite activity in motor control and learning, we organized our behavioral paradigm to induce a discrete change in movement trajectories under conditions known to require the premotor cortex (***Guo et al., 2014b***; ***Li et al., 2015***; ***Makino et al., 2017***; ***Bollu et al., 2021***; ***Dasgupta et al., 2025***). We monitored neural activity and tongue trajectories before (*i.e.,* pre-shift) and after (*i.e.,* post-shift) relocation of two lickport targets. Mice were trained to perform an auditory-cued directional licking task (Figure 1A; ***Guo et al., 2014b***; ***Inagaki et al., 2018***) with two target lickports approximately equidistant from the mouth (Figure 1A,B). Once the animals reached expert performance, we collected neural recordings and high-speed videography of tongue kinematics across multiple pre-shift behavioral sessions (Figure 1B; *N* = 22 mice, *N* = 4 median pre-shift sessions per animal, *N* = 306 median trials per session; see Methods). To extract a low-dimensional summary of each lick trajectory that was intuitive and relevant to the task outcome, we calculated the angle of the tongue tip as it exited the mouth at a threshold distance (Figure 1C). As expected, exit angles during correct trials formed two distinct clusters in the space of exit angle (Figure 1D). For each animal, we used linear discriminant analysis to define a choice boundary (CB) that distinguished right and left licks during pre-shift performance (Figure 1D; Figure 1—figure supplement 1A).

**Figure 1.**
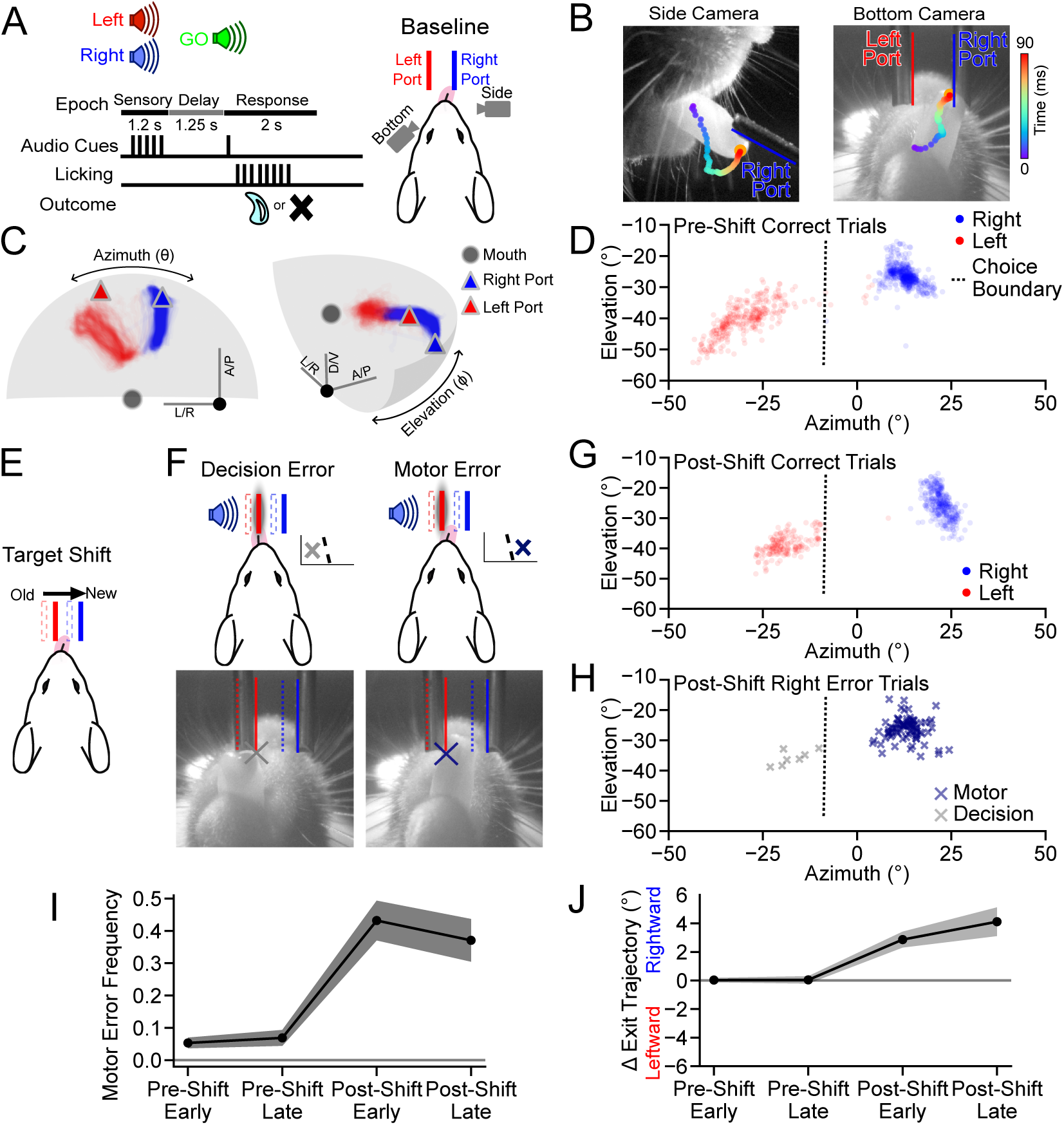
Mice learn new contingency-dependent movement trajectories after the relocation of targets. (A) Structure of the delayed-response directional licking task. (B) Single frames from the side and bottom cameras with the history of a tongue tip trajectory indicated by colored markers. (C) Example mouse pre-shift lick trajectories and lickport locations (Correct Right: blue, *N* = 539; Correct Left: red, *N* = 347 trials). Gray circle (“mouth”) indicates the intraoral reference point (*i.e.,* midpoint between the temporomandibular joints). Gray manifold is the distance threshold for lick exit angle calculation (D/V: dorsal/ventral; A/P: anterior/posterior; L/R: left/right). Trajectories only include time where the tongue tip is visible to both cameras. (D) Lick exit angles for all pre-shift Correct Right and Correct Left trials in (C). Dotted line denotes the choice boundary (CB). (E) Schematic of the port shift. Note: shifted left port location obstructs old rightward tongue trajectory. (F) Classification of right-cued error trials based on exit angle of the first lick with respect to the CB. (G) Lick exit angles for all post-shift correct trials for mouse in (D) (CR: Correct Right, *N* = 339; CL: Correct Left, *N* = 214 trials). (H) Lick exit angles for post-shift error trials from the mouse in (G) (MER: Motor Error Right, *N* = 87; DER: Decision Error Right, *N* = 7 trials). (I) Fraction of all right-cued trials that were motor error trials across training epochs (*N* = 22 pre-shift and *N* = 20 post-shift mice). (J) Relative lick angle (pre-shift normalized) for all first licks targeting the right port on right-cued trials. Shaded error and error bars: hierarchical bootstrap SEM. Bottom camera images are flipped to align with top-down views. **Figure 1—figure supplement 1.** Tongue exit angles change after the port shift to avoid the obstructing lickport.

To induce motor skill learning, we shifted the location of both lickports to the right (Figure 1E). 2P imaging and task timing, as well as contingencies between cue, choice, and reward, all remained unchanged after the shift (*N* = 20 mice, *N* = 4.5 median post-shift sessions per animal, *N* = 302 median trials per session). The new left lickport position encroached on space that had been occupied by the tongue during pre-shift licks towards the right port (Figure 1E,F). Thus, on right-cued trials, mice continuing to follow the pre-shift motor plan would be biased to more frequently contact the left port (Figure 1E,F; 30 ± 5% of pre-shift trajectories came within a tongue-width of the new location) and receive punishment (*i.e.*, timeout). We used the exit angle CB to distinguish this type of motor error – where the animal intended to lick the right port but contacted the left port –from decision errors, where the animal simply chose the left port (Figure 1F). Exit angles after the shift exhibited a bimodal distribution across both correct and error trials (Figure 1G,H; Figure 1— figure supplement 1C,D), supporting this distinction in error type. The frequency of licks classified as motor errors on right-cued trials increased significantly after the shift (median pre-shift 0.06, median post-shift 0.42, *p* < 0.001; Figure 1I; Figure 1—figure supplement 1C,D), reflecting the new challenge of avoiding the left lickport. In contrast to after the shift, exit angles on error trials before the shift were unimodal (Figure 1—figure supplement 1C). These errors were classified based on the fixed CB in order to demonstrate that very few pre-shift licks qualify as motor errors (Figure 1I), and not to suggest that tongue trajectories prior to the shift are behaviorally well-separated. Exit angles of licks directed towards the correct port (see Methods) shifted significantly rightward after the shift (late post-shift 4.11 ± 0.99^◦^, *p* < 0.001; Figure 1J; Figure 1—figure supplement 1), indicating that mice had learned a new contingency-dependent tongue trajectory.

### Longitudinal 2P calcium imaging in the apical tuft dendrites and somata of L5 ET neurons in ALM cortex

To monitor activity in L5 ET ALM neurons, we injected retrograde adeno-associated virus (AAV) encoding Cre recombinase into left ventromedial (VM) thalamus and AAV encoding Cre-dependent GCaMP8m (***Zhang et al., 2023***) into left ALM (Figure 2A). In *post hoc* histological sections, fluorescent neurons were restricted to left ALM cortex and no axonal labeling was detected in the corpus callosum (Figure 2B), indicating that our injections labeled L5 ET neurons and not intratelencephalic (IT) neurons. Daily imaging sessions alternated between L1 dendritic tuft and L5 somatic imaging (Figure 2D). To mitigate the effects of axial brain motion and capture more tuft dendrites at high imaging speeds (44.6 Hz frame rate), we implemented an excitation pulse splitting strategy that enabled extended depth-of-field (≈ 12 µm) 2P imaging in L1 (Figure 2E,F; Figure 2—figure supplement 1). Imaging of L5 somata was conducted with a standard 2P depth-of-field and interleaved line blanking (see Methods) to achieve high-fidelity imaging deep within the cortex at powers that were safe for longitudinal imaging (Figure 2F).

**Figure 2.**
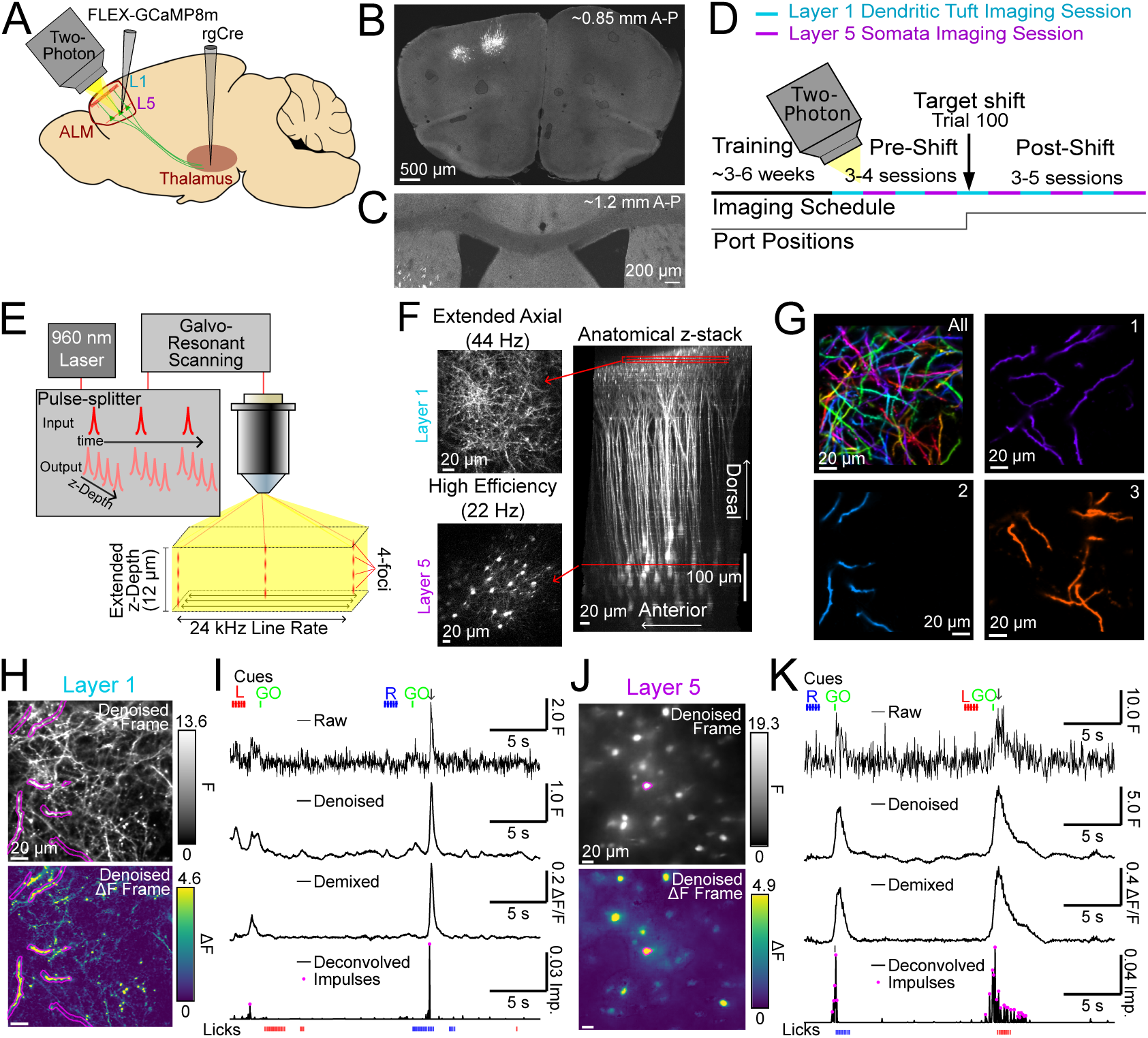
Longitudinal 2P calcium imaging of the tuft dendrites and somata of L5 ET neurons. (A) L5 ET neuron labeling approach. (B) Image of an example coronal section showing labeled neurons in left ALM cortex (∼0.85 mm along the A-P axis). (C) Image of the corpus callosum from a different section (∼1.2 mm A-P) of the same mouse as (B) showing the absence of axonal label. (D) Experimental timeline of dendritic and somatic imaging as well as the lickport shift. (E) Pulse splitting approach to obtain extended depth-of-field 2P imaging of L1 dendrites. (F) Mean projections (over a 10 s window) of functional imaging of L1 dendrites and L5 somata (left) along with their respective locations (red lines) in an x-*z* projection of an anatomical imaging volume (right). (G) Overlay of all L1 dendritic NMF spatial components from one animal (All; top left). Subsequent panels show three individual components. (H) Example denoised single frame from L1 functional imaging (top) and Δ*F* frame (bottom). Borders correspond to a binarized ROI derived from spatial component (2) in (G). (I) Example timecourses for ROI in (H) at different stages of processing: (i) raw motion-corrected, (ii) denoised (iii) NMF-demixed (iv) deconvolved. Pink dots denote impulse locations after filtering. Lines at the top indicate timing of the right cue (blue), left cue (red), GO cue (green) and the arrow indicates the time of the frame in (H). Ticks at bottom are right (blue) and left (red) licks. (J) Same as (H), but for example L5 functional imaging. (K) Same as (I), but for the soma in (J). **Figure 2—figure supplement 1.** Optical design of module for extended axial 2P imaging. **Figure 2—figure supplement 2.** Image registration, denoising, and segmentation pipeline. **Figure 2—figure supplement 3.** Estimated event kinetics, noise, and impulse rates across longitudinal 2P imaging.

To accurately track the activity of densely labeled dendritic arbors across many imaging sessions, we developed a custom image processing and segmentation pipeline that we used for both compartments (Figure 2—figure supplement 2A; see Methods). We used denoising (***Lecoq et al., 2021***; see Methods) and non-negative matrix factorization (NMF) to extract spatial components for each compartment (*i.e.* regions of interest; ROIs) from which we could isolate temporal components (*i.e.* traces) that minimized contamination from crossing or neighboring structures (Figure 2G; Figure 2—figure supplement 2A,E-F). Power spectral density of denoised traces and the coherence-weighted power spectral density of raw traces match up to approximately 2.6 Hz (Figure 2—figure supplement 2G; see Methods), which is not far from the bandwidth of GCaMP8m, given its estimated combined rise and decay (Figure 2—figure supplement 3B,C). To mitigate the impact of compartmental differences in calcium signal kinetics on downstream analyses, we then deconvolved (***Pnevmatikakis et al., 2016***) the temporal components into impulses (Figure 2H-K; Figure 2—figure supplement 3A-C; see Methods). Estimated measurement noise and impulse frequencies were stable across days in both dendrites and somata (Figure 2—figure supplement 3D,E), indicating that our imaging conditions did not cause systematic changes in brightness or overall activity. Impulse rates were significantly higher in somata compared to tuft dendrites (median dendrites: 0.138 Hz *N* = 402 ROIs, *N* = 21 animals; median somata: 0.240 Hz *N* = 528 ROIs, *N* = 20 animals; *p* < 0.05 K-S test), consistent with previous measurements from L5 neurons in the visual cortex (***Francioni et al., 2019***). Together, this longitudinal imaging paradigm provided us with a uniquely comprehensive readout of population activity in both L1 tuft dendrites and L5 ET somata across learning.

### Tuft activity tracks instructional cues more than planned action

Functional diversity across tuft dendrites has been observed in small numbers of L5 neurons in frontal cortex (***Cichon and Gan, 2015***; ***Kerlin et al., 2019***; ***Otor et al., 2022***). However, systematic differences in the sensorimotor selectivity of tuft dendrites and somata of these neurons have not yet been investigated. To address this, we first focused on analysis of data collected during the pre-shift period. Individual tuft dendrite ROIs (Figure 3A-C) and somata (Figure 3D-F) exhibited trial-type-specific activity that was generally consistent across sessions. To visualize task-related activity across the population of both dendrites and somata, we collected correct response trial types, calculated the mean GO-cue-aligned responses and sorted ROIs according to their trial-type preference and peak activity (Figure 3G,H; see Methods). Both tuft and somata populations exhibited diverse selectivity for trial type and a range of activity peaks that spanned the duration of trials. Upon close inspection of trial-to-trial activity, we observed that the onset of tuft activity was consistently time-locked to the GO cue (vertical green line; Figure 3B). This was in contrast to somatic activity, which had more variable timing (Figure 3E). We hypothesized that peri-GO tuft activity was more temporally aligned with the GO cue and that somatic activity was more aligned with the trial-to-trial timing of the initiation of action (Figure 3I,J). To systematically test this, we modeled the activity of each dendrite and soma as the sum of two nonnegative response functions: one response locked to the timing of the GO cue and another locked to the timing of first port contact (Figure 4A; see Methods). GO-associated activity sharply increased immediately after the GO cue in both dendrites and somata (Figure 4B,C; somata activity 2.58 ± 0.36, *p* < 0.001; dendrites 2.78 ± 0.24, *p* < 0.001; null hypothesis 1.0; between compartments *p* = 0.67), whereas contact-associated activity during this period was greater in somata than dendrites (Figure 4B,C; somata activity 2.02 ± 0.18, *p* < 0.001; dendrites 1.15 ± 0.13, *p* = 0.25; null hypothesis 1.0; between compartments *p* < 0.001) and not significantly different than baseline in dendrites. Thus, we found that peri-GO activity in the tufts is predominantly aligned with the GO-cue, whereas somata activity reflected both the timing of the GO-cue and the timing of action. To our knowledge, this is unique evidence of a strong functional distinction between tuft activity and somatic activity during motor behavior.

**Figure 3.**
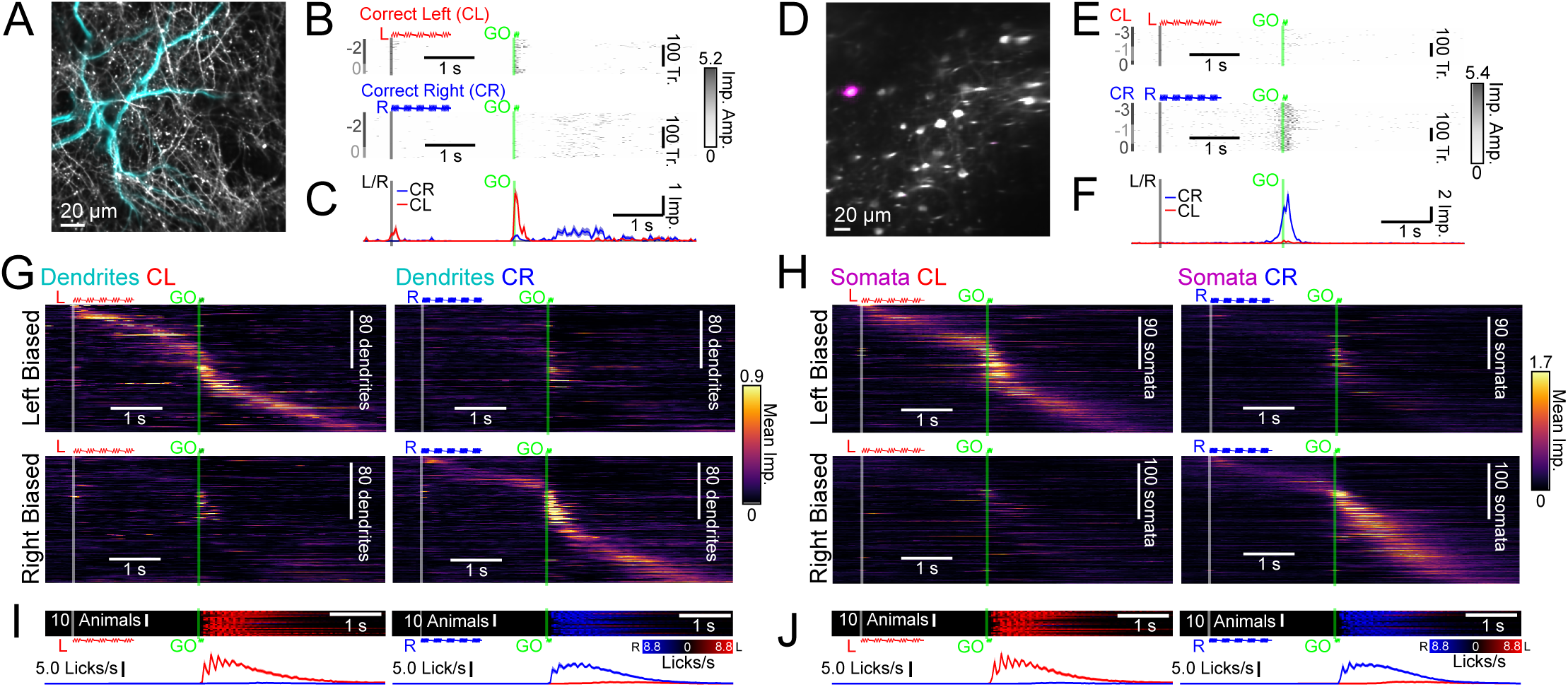
Diverse task-modulated activity in tuft dendrites and somata. (A) An example dendrite spatial component (cyan) overlaid on mean projection (gray). (B) Trial-aligned activity of the example in (A) for all pre-shift Correct Left (CL; *N* = 163) and Correct Right (CR; *N* = 212) trials. Session order identity of trials relative to the session of the lickport shift (session 0) is indicated by dark and light gray bars to the left of the rasters. (C) Mean trial-aligned activity for the dendrite in (A,B) on CR (blue) and CL (red) trials. (D-F) Same as (A-C), but for an example L5 soma (magenta) for all pre-shift Correct Left (CL; *N* = 282) and Correct Right (CR; *N* = 359) trials. (G) Mean pre-shift activity of dendrites for CL and CR trial types. ROIs are sorted by response bias and peak timing (left: *N* = 177 ROIs, *N* = 20 animals; right: *N* = 181 ROIs, *N* = 20 animals; see Methods for sorting details). (H) Same as (G), but for somata (left: *N* = 221 ROIs, *N* = 19 animals; right: *N* = 231 ROIs, *N* = 20 animals). (I) Mean individual animal pre-shift right (blue) and left (red) lickport contact rates (top) and mean across all animals (bottom panels, *N* = 21 animals) during dendrite sessions. (J) Same as (I), but for soma sessions (*N* = 20 animals). Auditory cues are indicated (left: red, right: blue, GO: green). Shaded error: bootstrap SEM.

**Figure 4.**
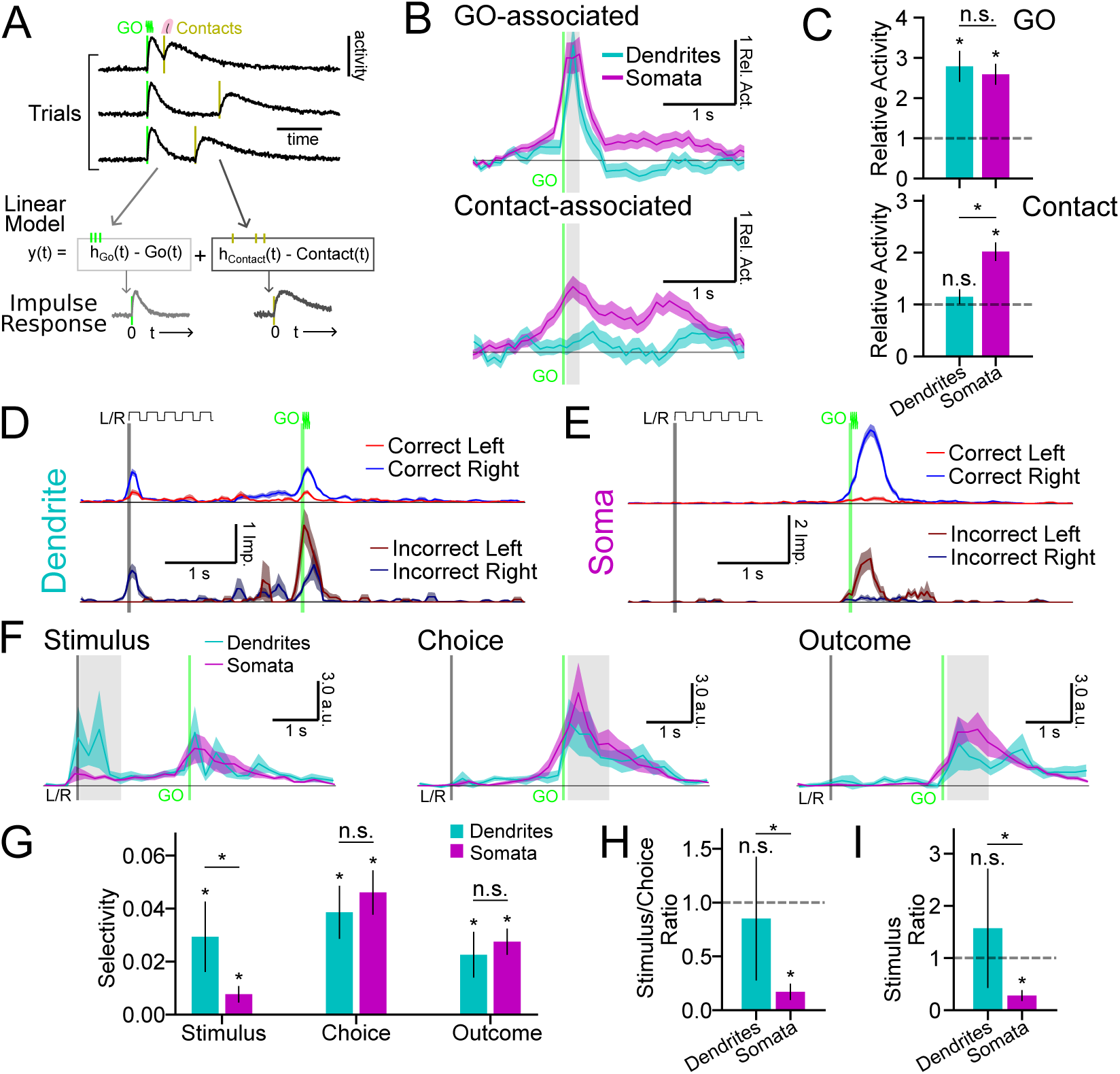
Tuft activity is less time-locked to action and more stimulus-selective. (A) Illustration of the estimation of GO-associated and contact-associated nonnegative response functions. (B) Mean GO-associated (top) and contact-associated (bottom) components of all dendrites (cyan) and all somata (magenta) relative to activity in the early delay period (see Methods). Contact-associated response functions were realigned to the timing of the GO cue before averaging. The vertical line indicates the start of the GO cue (green). (C) Mean relative activity during the shaded periods in (B). (D) Mean trial-aligned activity traces (pre-shift) for an example dendrite (CL: *N* = 264; CR: *N* = 334; IL: *N* = 15; IR: *N* = 74 trials). Sample cue (L/R; black) and GO cue (green) times are indicated. (E) Same as (D), but for an example soma (CL: *N* = 229; CR: *N* = 236; IL: *N* = 35; IR: *N* = 58 trials). (F) Mean normalized selectivity along the *Stimulus*, *Choice*, and *Outcome* CDs for dendrites (cyan) and somata (magenta) (see illustrated definitions in Figure 4—figure supplement 1A and Methods). Horizontal line: zero selectivity, left vertical line: sample cue start (L/R; black), right vertical line: GO cue start (green). (G) Mean normalized selectivity for *Stimulus*, *Choice*, and *Outcome* during the shaded time periods in (F). (H) Stimulus-selectivity to choice-selectivity ratio during the shaded time in (F). (I) Same as (H), but for stimulus-selectivity to outcome-selectivity ratio. Error bars: hierarchical bootstrap SEM. **Figure 4—figure supplement 1.** Mean trial-type projections along *Stimulus*, *Choice*, and *Outcome* CDs

To further understand the behavioral features encoded by activity in tuft dendrites and somata, we next considered activity during both correct trials and error trials (Figure 4D,E). Pre-shift, almost all errors were decision errors (decision error median 81.3% of all errors). Thus, behavioral trials could be categorized into four different trial types (Figure 4D,E) which differed in sensory instruction (*i.e.* initial cue), animal choice (*i.e.* port licked), and the task-outcome (*i.e.* presence or absence of reward). We modeled the mean activity of dendrite and soma ROIs during each trial type as a linear combination of these three task variables (***Chen et al., 2017***; ***Yang et al., 2022***). Since estimates of the selectivity of individual dendrites and somata tended to be noisy, we focused on population-level analyses of selectivity in an “activity space” where each dimension corresponded to the activity of individual dendrites or somata (***Stopfer et al., 2003***). From the linear model, we calculated coding directions (CDs) at each timepoint that maximally separated *Stimulus*, *Choice*, and *Outcome* activity (Figure 4F; Figure 4—figure supplement 1A) and estimated the direction and selectivity along each dimension across time (Figure 4—figure supplement 1B-E; see Methods). Allowing CDs to rotate in time (Figure 4—figure supplement 1B,D) – although unconventional – ensured that comparisons of population selectivity across the two compartments were not biased by the selection of an arbitrary CD time window. Tuft dendrites exhibited higher stimulus selectivity than somata during presentation of the initial auditory tones (Figure 4F,G; dendrites 0.029 ± 0.013; somata 0.008 ± 0.003; inter-compartment *p* < 0.05), and higher relative stimulus selectivity compared to choice and task-outcome selectivity during the response epoch (Figure 4G-I).

In summary, we found systematic differences in the baseline functional representations within the tuft dendrites and somata. Tuft activity was strongly stimulus-selective and aligned to the timing of instructional cues. Somatic activity also encoded these features, but the representation of the stimulus was weaker — and the representation of action timing was stronger — than in the tuft dendrites.

### Distinct encoding of corrective action in tuft activity

We next investigated whether activity in the tuft dendrites reflected motor corrections made during skill learning (*i.e.*, post-shift). Theories (***Körding and König, 2001***; ***Guerguiev et al., 2017***; ***Sacramento et al., 2017***) and recent experiments (***Francioni et al., 2026***) have suggested that apical dendrites are a critical locus of credit assignment calculations in the cortex. However, the nature of these calculations and their relationship with precise behavioral events remain unclear. In our motor learning paradigm, mice not only made behaviorally distinct errors across trials (Figure 1H), but also exhibited distinct behaviors after a motor error (Figure 5A). We focused our analysis on trials in which the exit angle of the first lick indicated intent to target the correct lickport (*i.e.*, based on the choice boundary; Figure 1H). These trials were then divided into three distinct trial types based on the success or failure of the first lick and the nature of any subsequent lick (Figure 5A). On Correct Right (CR) trials, the first lick made contact with the correct port and reward was delivered. On Correction Attempted (CA) trials, the first lick made contact with the incorrect port, and the mouse chose to direct a second lick toward the correct port (Figure 5A; prevalence: 54% of motor error trials). We interpreted these licks as a corrective action, because the tongue exit angle shifted further toward the correct target (Figure 5B). Abandoned Port (AP) trials were the same as CA trials, except the mouse either did not make a second attempt or the second lick was directed toward the incorrect port (Figure 5A; prevalence: 46% of motor error trials).

**Figure 5.**
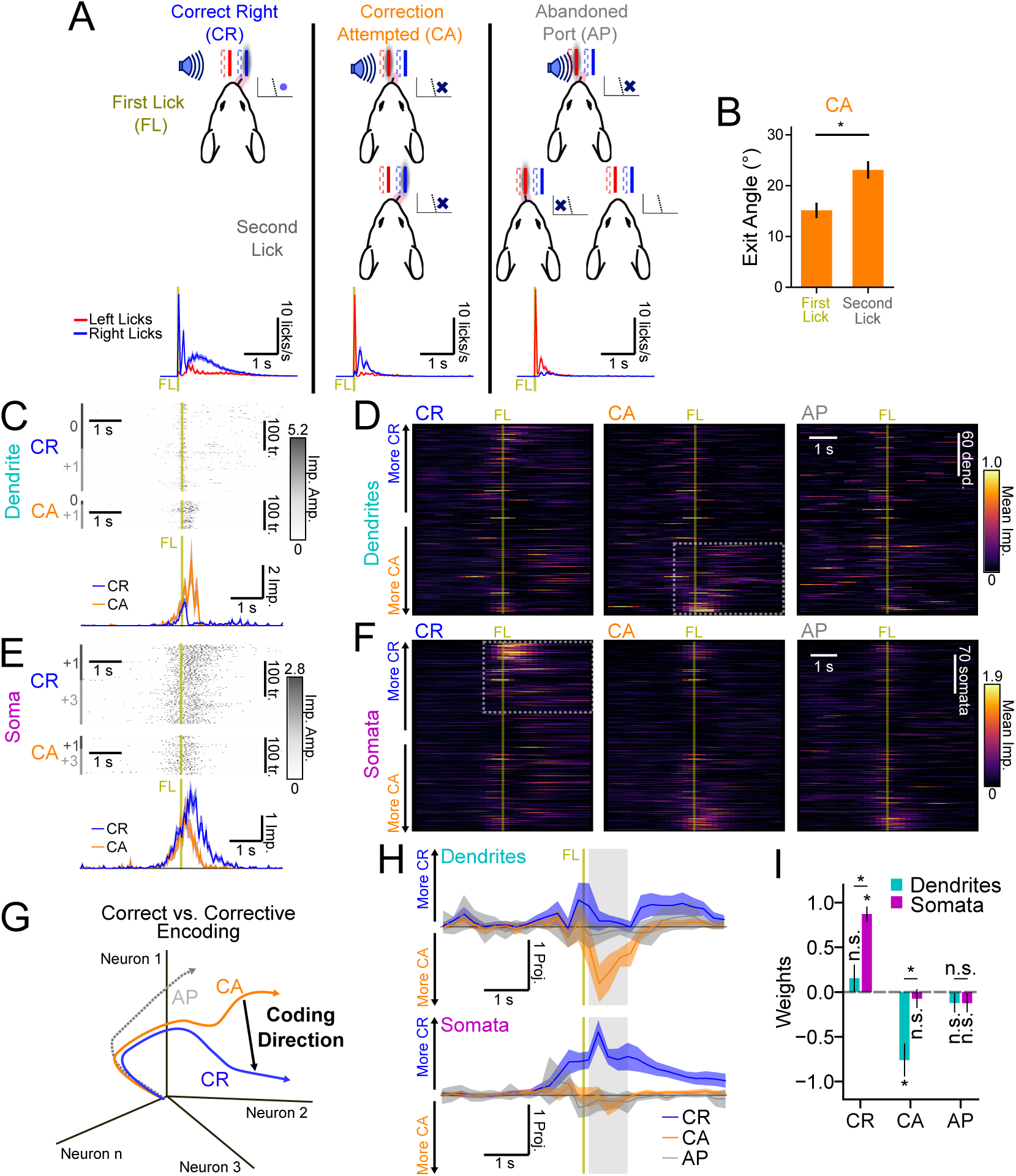
Distinct encoding of corrective action in tuft dendrite activity during motor learning. (A) Illustration of the classification of right-cued, right-choice trials based on the first lick (top row) and the second lick (middle row). Bottom row shows mean left (red) and right (blue) lickport contact rates aligned to the first lick (FL) contact (vertical dark yellow line) for all animals (*N* = 14). (B) Mean lick exit angle on CA trials for the first and second licks (*N* = 20 animals). (C) An example of dendrite activity on CR (*N* = 295) and CA (*N* = 94) trials (top) and mean first lick (FL)-aligned activity on CR (blue) and CA (orange) trials (bottom). The recording sessions (shift day 0) are indicated by alternating dark and light gray bars to the left of activity. (D) Mean activity of all recorded dendrites on CR, CA, and AP trial types (*N* = 261 dendrites, *N* = 14 animals; see Methods for inclusion criteria). Dendrites are sorted according to *CR* − *CA* activity between 0 and 1 s after the FL contact cue (see Figure 5—figure supplement 1). Dotted box highlights activity in CA-biased ROIs. (E) Same as (C), but for an example soma (CR: *N* = 225; CA: *N* = 109 trials). (F) Same as (D), but for all somata (*N* = 337 somata, *N* = 12 animals). The dotted box highlights activity in CR-biased ROIs. (G) Illustration of *CR* − *CA* coding direction in activity space. AP activity was independently projected onto the *CR* − *CA* CD. (H) Cross-validated projections of dendritic (top panel) and somatic (bottom panel) population activity onto the *CR* − *CA* CD for CR, CA, and AP trial types. The vertical line shows the time of first port contact while the horizontal line shows the zero projection. (I) Mean projection weight defined as the average projection normalized by the sum of CR and CA projections during the time denoted by the gray shaded area in (H). Shading and error bars: hierarchical bootstrap SEM. **Figure 5—figure supplement 1.** Differences in mean trial-aligned CR, CA, and AP activity. **Figure 5—figure supplement 2.** *CR* − *CA* code stability and additional trial-type projections.

Qualitative inspection of dendritic activity during these trial types revealed a sparse subset of dendrites exhibiting especially strong activation on CA trials after the first lick, but not on other trial types (Figure 5C,D; Figure 5—figure supplement 1). In contrast, the most strongly activated somata were selectively active on CR trials (Figure 5E,F; Figure 5—figure supplement 1). To quantify these differences, we again utilized population-level analyses. We focused on differences in activity between CR and CA trials, since neither compartment was particularly active on AP trials. We calculated a CD for each time bin that best distinguished CR and CA trials and then projected CR and CA trials along the CD (Figure 5G; Figure 5—figure supplement 2A). The projection of CA dendritic activity along this dimension was large after the first lick (Figure 5H,I; see Methods for normalization details). This was similar in magnitude to projection along a CD that distinguished CR and Correct Left (CL) trials (Figure 5—figure supplement 2B,C) – whereas projection of CR activity was comparatively weak. Projections of somatic activity along this dimension exhibited the opposite pattern (Figure 5H,I). The dendritic CA projection was significantly larger in magnitude than the somatic CA projection (Figure 5I; dendrites: −0.759 ± 0.18; somata: −0.074 ± 0.10; *p* < 0.01), while the somatic CR projection was significantly larger than the dendritic CR projection (dendrites: 0.175 ± 0.14; somata: 0.872 ± 0.08; *p* < 0.05).

The CD that distinguishes activity on CR trials and CA trials could reflect selectivity for many different behavioral variables that differ between the two trial-types. First, it could reflect simple selectivity for task-outcome (*i.e.*, reward or punishment). However, projections of activity on CL and Decision Error Right (DER) trials onto this CD were near zero for both dendritic and somatic activity (Figure 5—figure supplement 2C), indicating that the CD reflects selectivity for task-outcome that is specific to the contingency (*i.e.*, “lick right on right-cued trial”). To determine if activity was additionally selective for the presence or absence of corrective action, we projected AP activity along the same CD, since initial action and task outcome were the same on AP and CA trials. For somata, the projection of AP activity was similar to the projection of CA activity (Figure 5H,I; AP −0.123 ± 0.10, *p* = 0.115; CA −0.074 ± 0.10, *p* = 0.442), suggesting that this CD did not distinguish between follow-up actions in somata. However, for dendrites, the projection of AP activity was lower than the projection of CA activity (Figure 5H,I, AP −0.118 ± 0.10, *p* = 0.38; CA −0.759 ± 0.18, *p* < 0.05), indicating that dendritic activation along this dimension was selectively associated with corrective action. In summary, tuft dendrites encode a highly selective representation of corrective action, consistent with theories that posit a central role for tuft activity in credit assignment and learning.

### Divergent changes in tuft and somata function across motor skill learning

Learning can trigger transient and sustained reorganizations of cortical representations (***Costa et al., 2004***), including changes in the activity of the tuft dendrites of L5 neurons (***Lacefield et al., 2019***; ***Benezra et al., 2024***; ***Schoenfeld et al., 2021***). However, whether the tuft dendrites and somata of L5 neurons undergo similar or divergent functional changes during motor skill learning remains unknown. To address this, we compared the trial-aligned activity of individual tufts (Figure 6A) and somata (Figure 6B) before and after shifting the lickports. We focused our analysis on correct trials because the lick pattern (i.e., one contact lick followed by consummatory licking) on correct trials remained similar across the shift, facilitating clearer interpretations of any change in activity pattern.

**Figure 6.**
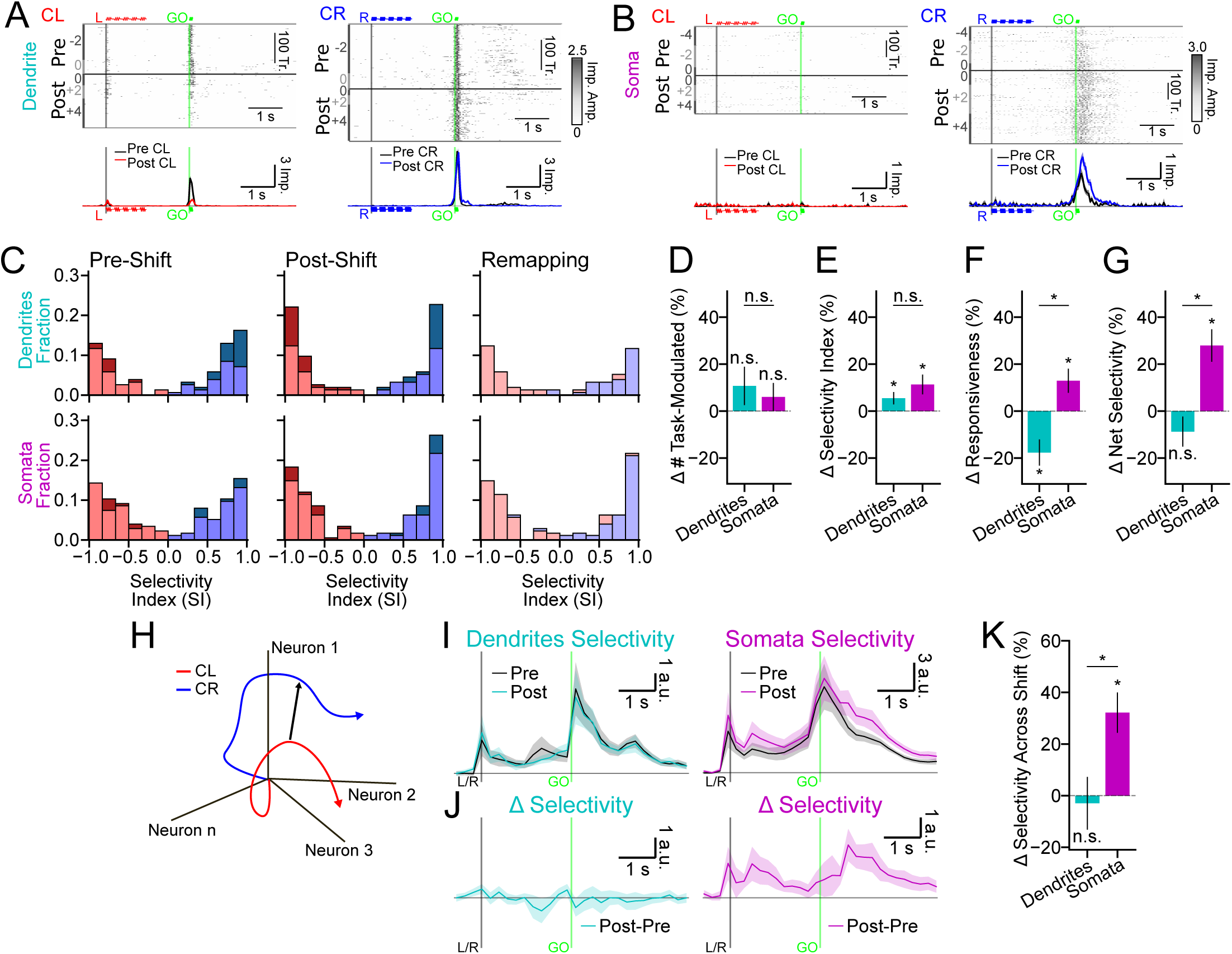
Compartmentalized changes in representation and gain across motor skill learning. (A) An example dendrite on CL trials (*N* = 163 pre-shift, *N* = 175 post-shift trials) and CR trials (*N* = 212 pre-shift, *N* = 170 post-shift trials) (top) and the trial-aligned mean activity (bottom) during the pre-shift (black) and post-shift training epochs (post-shift CL: red; post-shift CR: blue). Session order identity (session of shift = 0) of the trials is indicated by dark and light gray bars to the left of the rasters. Auditory cue timings indicated. (B) Same as (A), but for an example soma on CL (*N* = 288 pre-shift, *N* = 223 post-shift trials) and CR (*N* = 258 pre-shift, *N* = 432 post-shift trials) trials. (C) Selectivity index (SI) distributions for task-modulated ROIs (see Methods). Pre-shift (left column) and post-shift (middle column) show SI distributions for dendrites (top) and somata (bottom). *Lighter* colors are proportion in each bin that were task-modulated in both training epochs, whereas *darker* colors are the proportion uniquely task-modulated in that training epoch. Remapping (right column) only shows ROIs that were task-modulated in both training epochs (red portions: preferred left during pre-shift; blue portions: preferred right during pre-shift). (D) Percent change in the number of task-modulated dendrites and somata across the port shift. (E) Percent change in mean selectivity index ([*CR* − *CL*]∕[*CR* + *CL*]). (F) Percent change in mean responsiveness (*CR* + *CL*). (G) Percent change in mean selectivity magnitude (*CR* − *CL*). (H) Illustration of the *CR* − *CL* CD in activity space. (I) Cross-validated selectivity between CR and CL projections along the *CR* − *CL* CD for dendritic (left) and somatic (right) ROIs during the pre-shift (black) and post-shift (colored) training epochs. Vertical lines indicate start of the sample (black) or GO (green) cue. (J) Difference between pre-shift and post-shift selectivity projections. (K) Percent change in selectivity (I) from pre-shift to post-shift, averaged across the whole trial period. Shading and error bars: hierarchical bootstrap SEM. **Figure 6—figure supplement 1.** Trial-type projections before and after the lickport shift. **Figure 6—figure supplement 2.** Selectivity CD magnitude across early and late learning. **Figure 6—figure supplement 3.** Selectivity CD correlations across early and late learning.

Learning can be accompanied by many different changes in cortical representations, such as changes in the prevalence, selectivity, and responsiveness (*i.e.*, gain) of feature-sensitive neurons. To distinguish between these options, we first identified the ≈0.5 s time window with the largest difference in activity between CR and CL trials for each dendrite or soma (see Methods). In this time window, we then calculated a selectivity index (SI) in each training epoch (*i.e.*, pre- or post-shift) as the mean difference between CR and CL responses divided by their sum. If the SI was reliable (95% CI < 0.66), the dendrite or soma was considered “task-modulated” in that training epoch regardless of the SI value. In both dendrites and somata, the post-shift distribution of SI shifted towards the edges compared to pre-shift (Figure 6C). A very small fraction of dendrites or somata changed preference across the shift (Figure 6C; dendrites *N* = 5 or 3%; somata *N* = 7 or 4%), indicating minimal population remapping. The prevalence of task-modulated dendrites and somata also did not significantly change across the shift (Figure 6D; Δ selective dendrites 11.98 ± 8.08%, *p* = 0.065; somata 6.78 ± 5.75%, *p* = 0.212). However, SI values in both compartments increased (Figure 6E, Δ SI dendrites 5.43±2.15%, *p* < 0.05; somata 11.16±3.73%, *p* < 0.01). Surprisingly, the port shift induced divergent changes in the responsiveness of both compartments (Figure 6F; dendrites −17.75 ± 5.01%, *p* < 0.001; somata 12.06 ± 4.52%, *p* < 0.01; between compartments *p* < 0.001). Dendritic responses decreased, whereas somatic responses increased. The overall impulse rates of dendrites and somata during behavior remained steady across the shift (Figure 2—figure supplement 3E), so these changes were presumably specific to trial-type selective activity. We then calculated net selectivity as the difference between CR and CL responses (*i.e.*, without normalization by the sum). The net selectivity of dendrites did not change significantly across the shift (Figure 6G; −8.88 ± 6.04%, *p* = 0.127), whereas the net selectivity of somata increased (Figure 6G; 27.05 ± 6.04%, *p* < 0.001; between compartments *p* < 0.001).

To measure net selectivity across the population without any filtering for task-modulation, we again returned to population-level analyses in activity space. Within each training epoch, we calculated the CD that best distinguished activity between CR and CL trials (Figure 6H), cross-validated projections of CR and CL activity along these CDs (Figure 6—figure supplement 1A,B), and determined the selectivity projection as the difference between these projections (Figure 6I). The difference between the pre- and post-shift selectivity (Figure 6J) confirmed that net selectivity did not change across training epochs in the tuft dendrites, but increased substantially in the somata (Figure 6K; dendrites −3.08 ± 9.89%, *p* = 0.724; somata 32.18 ± 7.60%, *p* < 0.001; between compartments *p* < 0.01). This increase in the magnitude of the net selectivity of somata occurred early in learning (first 2 training sessions after the shift) and then returned to near-baseline in later sessions (Figure 6—figure supplement 2), consistent with previous studies of L5 neuron output in frontal cortex during learning (***Costa et al., 2004***; ***Peters et al., 2017***; ***Hwang et al., 2021***). Changes across learning in the direction of the CD were not significantly different between dendrites and somata (Figure 6—figure supplement 3), consistent with their similar magnitude of changes in SI across the shift (Figure 6E).

In summary, the tuft dendrites and somata exhibited divergent changes in the gain of task-modulated responses across learning. Increases in dendrite selectivity were tightly balanced by decreases in gain, suggesting that tuft activity is regulated to maintain consistent net selectivity across learning. In contrast, the net selectivity of the somata increased transiently during early learning.

## Discussion

Previous work identified diverse behavior-related dendritic activity that was localized to different compartments within individual layer 5 neurons in the frontal cortex (***Kerlin et al., 2019***; ***Otor et al., 2022***). However, any systematic functional differences between regenerative activity in the tuft dendrites and somatic output remained unclear. In the present study, we uncovered systematic differences between the tuft dendrites and somata in sensorimotor selectivity, dynamics, corrective signaling, and response gain across different phases of motor learning. These differences in encoding and functional plasticity suggest specialized roles for tuft computation in motor control and motor learning, and provide a critical new foundation for further investigation of the contributions of dendritic computation to behavior.

### Selectivity and timing of L5 tuft activity during expert behavior

We found that, during the sensory cue period of the licking task, tuft dendrites of L5 neurons in ALM cortex are more stimulus-selective than somata. This is consistent with the general role for tuft spikes in conditioning the activity of L5 somata on feedback signals (***Larkum, 2013***; ***Xu et al., 2012***; ***Manita et al., 2015***; ***Ranganathan et al., 2018***; ***Fişek et al., 2023***; ***Zhang et al., 2025***) and recent work indicating that suppression of L1 activity in ALM cortex prevents relearning of context-dependent rules (***Maristany de las Casas et al., 2026***).

More surprisingly, we found that tuft activity in ALM cortex exhibited distinct timing that was associated with the GO cue, but not the trial-to-trial timing of action initiation. In contrast, somatic activity was associated with both GO cue and action initiation. Interestingly, ventromedial (VM) motor thalamus provides stronger input to the apical tuft than to the basal dendrites of Layer 5 ET neurons in ALM (***Guo et al., 2018***). Cue-triggered VM input rapidly shifts ALM activity along a distinct “GO” mode prior to movement onset (***Inagaki et al., 2022b***). However, this shift along the “GO” mode is not sufficient to initiate movement. Instructive activity along a different activity mode, dominated by the output activity of L5 ET neurons, prepares and executes directed movement (***Li et al., 2015***; ***Inagaki et al., 2022b***). Our finding that tuft activity in L5 ET neurons in ALM is predominantly aligned to the GO cue raises the possibility that it could drive or reflect the transition along the “GO” mode. Tuft spikes could drive bAP-activated calcium (BAC) burst firing (***Larkum et al., 1999b***), signaling the state transition within a movement-null space (***Kaufman et al., 2014***) to downstream structures, as well as back to L5 IT and ET neurons in ALM *via* low-latency loops (***Guo et al., 2018***). Subsequent somatic firing, uncoupled from the tuft, could control action within a movement-potent space. Consistent with these ideas, recent models posit that sparsely active (***Logiaco et al., 2021***) or weakly selective (***Pereira-Obilinovic et al., 2025***) thalamic inputs can rapidly transition the cortical network towards particular future actions.

### Representation of corrective action in the tuft of L5 ET neurons

Frontal cortex is a critical node in an action control system that selectively minimizes goal-relevant errors (***Todorov and Jordan, 2002***; ***Scott, 2012***) through both prospective internal (***Kao et al., 2021***) and ongoing sensory (***Sauerbrei et al., 2020***; ***Kalidindi et al., 2021***) feedback control. Information regarding prediction errors must be utilized to generate real-time corrections and to remodel the context-specific networks that generated the action in order to improve future movement. How corrective action is represented in the cortex is still debated (***Perich et al., 2024***; ***Ames et al., 2019***; ***Kudryashova et al., 2025***; ***Malonis et al., 2021***) and circuit mechanisms linking corrective action to the plasticity of individual neurons remain unclear.

We found that strong activation of the tuft population was selectively associated with corrective action, while the somatic population was particularly active during reward and generally active during action (*i.e.*, regardless of whether that action was corrective or not). The deconvolved tuft signal associated with a corrective lick peaked ≈200 ms after the average time of second port contact and remained elevated for another ≈500 ms. This is longer than GCaMP8m kinetics (Figure 2—figure supplement 3A-C; ***Zhang et al., 2023***), suggesting that most of the tuft signal occurred after the initiation of the corrective movement. Thus, our results favor a model in which the tuft spikes associated with corrective action are involved in post-correction computations (*e.g.*, driving plasticity) rather than driving the corrective action itself. Our finding that strong tuft activity selectively occurred during corrective action suggests that nonlinear integration of error (*i.e.*, “target missed”), context (*i.e.*, “right port is the target”), and movement (*i.e.*, “second lick is rightward”) occurs either within the tuft or upstream of it. A recent study in retrosplenial cortex found that dendritic spikes in neurons were modulated by performance signals related to the causal role of each neuron in a neurofeedback-based task (***Francioni et al., 2026***). Although we do not know whether the strong correction-related activity we observed played a causal role in learning, the specificity of the behavioral circumstances that generated this signal could be consistent with a responsibility-weighted error signal to the subnetwork of L5 ET neurons that controlled the contingency-specific action (*i.e.*, rightward choice). If this correction-associated signal, observed in the tuft dendrites of L5 ET neurons, indeed reflects a credit assignment calculation, many afferent inputs to the tuft could provide the necessary information for such a calculation, including the prediction error (***Manita et al., 2015***; ***Ako et al., 2025***; ***Heindorf et al., 2018***; ***Levy et al., 2020***), the context-specific goal (***Moberg and Takahashi, 2022***; ***Inagaki et al., 2022a***), and an efference copy of past output (***Guo et al., 2018***). Now that this correction-associated tuft signal has been identified, future work can determine the precise circuit mechanisms generating it, as well as determine whether it is indeed a teaching-related signal or reflects some other correction-associated computation.

### Motor skill learning triggers compartmentalized changes in representation and gain

Our results demonstrate that across motor skill learning, the selectivity of tuft dendrites of L5 ET neurons in ALM cortex increases. This is similar to the enhanced selectivity for behaviorally relevant task dimensions after learning that has been observed in the tuft dendrites of L5 neurons in the somatosensory cortex (***Lacefield et al., 2019***; ***Benezra et al., 2024***; ***Schoenfeld et al., 2021***). Our recordings from the somata of L5 ET neurons in premotor cortex revealed a similar increase in selectivity. However, we found that motor skill learning was accompanied by a decrease in the response gain of task-modulated dendrites and an increase in the gain of task-modulated somata. To our knowledge, these divergent changes in the gain and net selectivity of L5 tuft dendrites and somata of frontal cortex have not been previously reported. The increase in the somatic net selectivity declined later in learning, consistent with previous findings in motor cortex (***Peters et al., 2017***).

Most surprisingly, the increase in selectivity of selective tuft dendrites was accompanied by a matched decrease in overall responsiveness, such that net selectivity remained stable across learning. This suggests that, even within a particular behavioral context, total dendritic activity and the sparseness of that activity may be tightly regulated. Interestingly, a recent study of L5 neurons of the auditory cortex found decreased, but more synchronous, activity across tufts during recall of conditioned stimuli and associated this change with an increase in *I*_ℎ_-current (***Rosier et al., 2025***). L5 neurons in visual cortex also exhibit divergent compartmental changes in activity following passive exposure to pattern-violating stimuli (***Gillon et al., 2024***).

We did not investigate causality, so the precise relationship between the neural changes we observe during learning and the behavior of the animal remains unclear. Given the port shift direction relative to the hemisphere we imaged, the increased net selectivity we observe could be directing licking trajectories towards the new port locations. Alternatively, the changes in net selectivity could reflect increased engagement of ALM cortex due to increased uncertainty about upcoming action (***Dragoi et al., 2026***). Since total activity did not change, the specific increase in the gain of neurons selective for the task could be considered a form of “motor attention” (***Dragoi et al., 2026***). This could also explain the decrease in the gain of the tuft dendrites, as some pathways that are more active during increased arousal or attention suppress the apical dendrites while simultaneously increasing the gain of the basal compartment (***Malina et al., 2021***).

### Study limitations

We refer to the behavioral paradigm as motor “skill learning” strictly based on the nature of the task. After the shift, mice must avoid a new obstacle close to the mouth (*i.e.,* the left port), which we assume requires the formation of a distinct motor controller memory and therefore would be considered skill learning (***Krakauer et al., 2019***). However, we did not confirm that the mice exhibited specific behavioral characteristics of skill learning and it is possible that other kinds of motor learning (*e.g.,* motor adaptation) were dominant.

L5 ET neurons in premotor cortex elaborate extensive tuft dendrites in L1, whereas Layer 6 (L6) corticothalamic (CT) neurons are predominantly untufted (***Jiang et al., 2020***; ***Peng et al., 2021***). Thus, although we cannot rule out the possibility that dendrites from a subclass of L6 CT neurons were also sampled, it is likely that the vast majority of dendrites we recorded in L1 originated from L5 ET neurons.

In order to detect sparse representations in the tuft, we adopted a dense labeling strategy that precluded unambiguous tracing of dendrites. The dendritic signals we measured could reflect bAPs and regenerative events that generate global, hemi-tree, or local branch calcium influx (***Hill et al., 2013***; ***Otor et al., 2022***). Based on previous calcium imaging of L5 tufts in ALM cortex of mice engaged in similar tasks (***Kerlin et al., 2019***; ***Maristany de las Casas et al., 2026***), we suspect that most of the activity we measured was coincident with global tuft or hemi-tree events, as well as somatic spiking. Recent *in vivo* voltage imaging in the hippocampus has also indicated that most spikes in distal dendrites start as bAPs that have been selectively amplified (***Lee et al., 2026***; ***Wu et al., 2026***). Future studies that directly manipulate dendrite conductances could begin to identify precise mechanisms and investigate causal relationships with behavior.

## Methods

All experiments were conducted in accordance with the NIH Guide for the Care and Use of Laboratory Animals and were approved by the Institutional Animal Care and Use Committee of the University of Minnesota. Data are from 22 mice (C57BL/6J, Jackson Laboratories, stock #000664) of either sex (82 ± 20 days old at the beginning of experiments). Eleven were male, and eleven female. There were no sex-specific differences in the results and data were pooled across sex.

### Surgery and Virus Injection

Cranial window surgeries were conducted as previously described (***Daie et al., 2023***). We injected viral retrograde Cre recombinase (pENN-AAV-hSyn-Cre-WPRE-hGH, Addgene; 1.9×10^12^ vg∕mL; 200 nL) through a burr hole in the skull into left thalamus (−1.5 mm posterior, 0.8 mm lateral, 3.8 mm deep relative to bregma). We next performed a 3 mm craniotomy over left ALM (2.5 mm anterior, 1.5 mm lateral relative to bregma) and injected the exposed cortex with FLEX-GCaMP8m (pGP-AAV-syn-FLEX-jGCaMP8m-WPRE, Addgene; 1.7 × 10^12^ vg∕mL; 75 nL per injection; 9 injections total in a 3×3 grid; depth of 700 µm below pia). A triple coverglass window (#1 coverglass, 2.5 mm / 2.5 mm / 3 mm; Potomac Photonics and Warner Instruments) covering the cortex and a headbar were affixed to the skull with adhesive (C&B Metabond; Parkell) and dental acrylic (Jet; Lang Dental).

### Behavioral Paradigm

During training and recording, mice were placed on water restriction (1 mL per day). Initial training and task structure were similar to previous studies (***Guo et al., 2014a***; ***Guo et al., 2014b***; ***Inagaki et al., 2018***) and shown in Figure 1A. All task epochs and triggers were controlled by a BPOD State Machine (Sanworks). At the beginning of each trial, mice were presented with an auditory sequence (5 tones of 0.15 s with 0.1 s between tones; total duration 1.2 s) of 3 kHz (left trial) or 18 kHz (right trial) pure tones. After presentation of these “sample period” tones, mice had to withhold licking for a 1.25 s delay until the presentation of an auditory GO cue (carrier frequency: 6 kHz, modulating frequency: 360 Hz) lasting 0.1 s. Port contact prior to the GO cue reset the timer for that epoch and these trials with “early” licking were excluded from further analysis. Mice had 2 s to respond by licking one of two lickports with internal edges (the typical locations of the first tongue contact) separated by a distance of 2 mm. Tongue contact with the correct port first (as reported by a custom electrical lick detector) resulted in a water reward (3 µL) at that port. Contact with the incorrect port first resulted in a 15 s timeout. Trials in which the mouse failed to respond after the GO cue were also excluded from further analysis. To ensure the animals learned a stereotyped licking behavior the lickports were fixed in position with respect to their head across pre-shift training sessions. Mice typically learned this task over the course of ≈25 training sessions. Once task performance reached threshold (>75% correct), baseline tongue trajectory and neural data (see Methods) was collected over ≈4 sessions. After recording baseline sessions, both lickports were shifted by 1 mm to the right of the mouse on the hundredth trial of the “shift” session (session 0). Across all sessions, all trials prior to the trial of lickport shift were considered “pre-shift” and all trials subsequent to that trial were considered “post-shift” for subsequent analysis. For more detailed analyses of learning dynamics, we specified 4 phases of learning: two phases before the shift (pre-shift early: >1 session before the session of the shift; pre-shift late: the session immediately prior and session of the shift up to the shift trial) and two phases after the shift (post-shift early: session of the shift after the shift trial and the immediately subsequent session; post-shift late: >1 session and less than 3 sessions after the shift session).

### Behavior Analysis

#### Tongue Tracking

Two CMOS cameras (FLIR Blackfly S, BFS-U3-04S2M) were located to the side and below the mouse (Figure 1A,B). Custom software (***Tran, 2026***) captured timestamped video at 500 Hz from both cameras for 2 s after every GO cue. The animal was illuminated with infrared LEDs. For analysis of the tongue trajectory, we first collected all recorded images from both the side and bottom cameras across all sessions and all animals. A custom convolutional neural network (***Dinh, 2026***) determined whether the tongue was visible outside of the mouth in each frame. To extract the tongue position in 3D, we then trained a second neural network using DeepLabCut (***Mathis et al., 2018***) on a subset of frames in which the tongue was visible. This network was used to estimate the tongue’s 3D coordinates for all frames containing a visible tongue. Using these outputs, we constructed a time-resolved trajectory for each lick synchronized with electrical lickport signals to determine if and when each lick contacted a lickport.

#### Lick Angle Quantification and Classification

Since jaw movements varied from mouse-to-mouse and trial-to-trial, an intraoral reference point (“mouth”) was defined as the approximate midpoint between the temporomandibular joints and used for tongue trajectory calculations. To reduce each 3D tongue trajectory to an intuitive estimate of movement towards the lickports, we calculated the exit angle (i.e., azimuth *θ* and elevation *ϕ*) of the tongue as it extended beyond a critical distance from the “mouth”. This distance was the minimum distance from the mouth to the closest lickport position across both pre-shift and post-shift lickport positions. Linear Discriminant Analysis (LDA; python scikit-learn; ***Pedregosa et al., 2011***) of pre-shift exit angles on correct trials was used to determine a choice boundary. Exit angle with respect to this boundary was used to define the intended target of the lick, regardless of the subsequent port contact.

Exit angle distance from the choice boundary (Figure 1—figure supplement 1) was calculated as follows. First, the lick coordinates (*θ*, *ϕ*) were projected onto the LDA decision line in 2D angle space. Then, each lick and its projection were converted into 3D unit vectors (**v**_lick_, **v**_proj_), originating from the animal’s mouth. We defined the lick angle as the angular separation (*ω*) between these vectors:

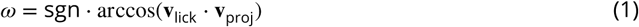

where sgn is the sign of the LDA decision function, assigning positive or negative values based on the classified lick target (positive = right, negative = left). For population analyses, we normalized each animal’s lick angle by subtracting the average lick angle calculated from the entire pre-shift period. This relative lick angle was then averaged across all trials within each training epoch, and then averaged across animals. For SEM estimates and hypothesis tests, we bootstrapped this process 10000× by drawing animals with replacement.

#### Gaussian Mixture Model Analysis

To assess whether the lick angle distributions were unimodal or bimodal, we fit one- and two-component Gaussian mixture models (GMMs) to the lick angle data using maximum likelihood estimation. Models were compared using the Bayesian Information Criterion (BIC). The difference in BIC between the one- and two-component models (ΔBIC = BIC_1_ − BIC_2_) was used to assess relative model fit, with positive values indicating evidence in favor of the two-component model.

### Functional 2P calcium imaging

For high-speed extended depth-of-field imaging of L1 dendritic tufts, we built a custom 2P pulse splitter system for multi-foci simultaneous excitation combined with high-speed scanning. This extended-depth-of-field 2P imaging approach maintained a high excitation efficiency by confining excitation light to four diffraction-limited foci. This allowed us to use average infrared laser powers that were suitable for long-term imaging. In brief, excitation light (960 nm; 100 fs; 80 MHz; InsightX3, Spectra Physics) was temporally multiplexed using a custom pulse splitter consisting of a series of polarization gated delay lines that produced an output pulse train of 320 MHz (see Figure 2E; Figure 2—figure supplement 1). Optical relays within each delay line imparted a variable divergence, resulting in the generation of four foci equally separated in time and space that spanned an approximate depth of 12 µm. For more details, see Figure 2—figure supplement 1. L1 imaging was conducted with a 25× microscope objective (Olympus XLPLN25XWMP2 25× NA 1.05) and average laser power exiting the objective of between 20–50 mW (mean: 34 mW, 0.11 nJ pulse energy per focus). The frame rate was 44.6 Hz and the field of view (FOV) was 210 µm × 210 µm × ≈ 12 µm.

A different configuration was used to achieve high signal-to-noise ratio recordings from somata in L5 at reasonable average laser powers. We blocked 80 MHz excitation light from reaching the sample on 3 out of every 4 imaging lines scanned by our resonant mirror. This was a cost-effective alternative to maintaining a separate, lower repetition rate light source. L5 imaging was conducted with a 16× microscope objective (Nikon CFI75 LWD, NA 0.8) and average laser power exiting the objective (with 75% line blanking) was 9 mW to 38 mW (mean: 22 mW, 1.1 nJ pulse energy). The frame rate was 22.8 Hz and the FOV was 328 µm × 328 µm.

In both imaging configurations, lateral scanning was performed by a resonant mirror (CRS 12 kHz, Novanta Photonics) conjugated to a set of galvanometer mirrors (6215H 5mm, Novanta Photonics). Emission photons were detected using a photomultiplier tube (PMT; PMT2101, Thor-labs), digitized (NI-5771/NI-7975R/PXIe-1092, National Instruments) and recorded with ScanImage (MBF Biosciences).

Somatic and dendritic imaging sessions alternated every other day (19 of 22 mice), except for 3 mice in which only one compartment was imaged daily (dendrite-only: 2 mice, soma-only: 1 mouse). These exceptions were due to brain curvature or the angle of the coverslip with respect to the brain, such that only one compartment could be imaged and the other compartment was underneath skull regrowth or dural thickening that made high-quality imaging impossible. For Figure 5 and Figure 6, which make comparisons between dendritic and somatic activity during the post-shift period, 67% of animals providing somatic data (4 of 6 mice) underwent somatic imaging on the day of the shift and 80% of mice providing dendritic data (8 of 10 mice) underwent dendritic imaging on the day of the shift.

### Anatomical 2P imaging

After completion of functional imaging, a *z*-stack was collected (328 µm×328 µm FOV; 44.6 Hz frame rate; 100 frames per step; 2 µm steps) from pia to L5 using a 16× microscope objective (Nikon CFI75 LWD, NA 0.8). Frames were registered within each slice, mean-projected within slice, aligned to neighboring slices, and then mean-projected along the stack *y*-axis.

### Histology

After completion of all data collection, animals were deeply anesthetized with ketamine-xylazine and a transcardiac perfusion was performed *via* the left ventricle with saline followed by 1:10 buffered formalin. The brain was extracted and post-fixed in formalin overnight at 4 °C, then rinsed in phosphate-buffered saline (PBS). Coronal sections of 60–100 µm thickness were collected with a vibratome and mounted in Vectashield (Vector Laboratories). GCaMP8m fluorescence was imaged (excitation wavelength: 488 nm) using an upright Nikon Ti2 confocal microscope with a Plan APO 10× NA 0.45 air objective. Whole coronal sections were imaged using tiles to cover the entire slice, stitched using NIS-Element (Nikon Imaging), and finally aligned to the Allen Brain Atlas (***Wang et al., 2020***). Histology images were spatially smoothed for display purposes (Gaussian *σ* = 2 µm; Figure 2B,C)

### Image Analysis

In order to extract activity traces for dendritic tufts and somata, image series underwent a combination of rigid registration, denoising, non-rigid warp correction, segmentation and finally deconvolution (Figure 2—figure supplement 2A). Front-end data intake and metadata management was performed using custom Matlab (Mathworks) scripts. All other imaging data were processed using custom Python scripts.

#### Rigid registration

All image series were subjected to an initial rigid registration (phase cross-correlation; no upsam-pling; python scikit-image; ***Walt et al., 2014***). All imaging from the same animal and same layer was registered to a common reference target. Offsets were estimated from spatially filtered (Gaussian *σ* ≤ 1 µm) frames, adaptively filtered to suppress registration noise, and then used to transform unfiltered frames.

#### Denoising

Following rigid registration, image series were denoised using DeepInterpolation (***Lecoq et al., 2021***), modified to minimize L2 (rather than L1) loss and to receive as its input, 10 mean projections of frames collected from non-overlapping exponential (base 1.75; exponents: 0–9) frame windows both preceding and following the interpolated frame. The DeepInterpolation model was trained on each imaging session prior to denoising of that session. Denoising prior to NMF-based segmentation resulted in more robust and consistent dendrite segmentation than NMF-based segmentation without prior denoising (0.79 ± 0.01 *ρ* vs. 0.46 ± 0.01 *ρ*; mean of the max Spearman correlation of components across sessions; random subsample of N = 3 mice, 15 sessions, 400 components).

For Figure 2—figure supplement 2G, power spectral densities (PSDs) were calculated for timecourses extracted using a fixed set of binary dendrite masks for each FOV (random sample: N=3 mice, 3 sessions, 161 dendrites) for both raw and denoised image series. Image series were pixel-wise corrected for brain motion, but not warp corrected, to avoid smearing measurement shot noise across pixels. To estimate the “latent” PSD of signals in the absence of shot noise, each ROI’s pixels were randomly split into two halves (i.e., replicates), traces extracted, and then the cross-spectral density was calculated using a segmented multitaper method. To estimate the “recoverable” PSD of latent signals given the shot noise and the duration of recordings, we calculated the coherence of the ROI replicate traces and then used these values to calculate the coherence-weighted PSD.

#### Non-rigid registration

Our segmentation methods (see Methods) required that all dendritic imaging from the same animal be very precisely aligned. Small within-session and across-session instabilities (*e.g.*, resonant mirror temperature changes, vessel dilation) resulted in image warp that was significant for longitudinal dendritic imaging. Warp calculations were performed on contrast-enhanced frames using symmetric diffeomorphic registration (python DIPY; ***Garyfallidis et al., 2014***; ***Avants et al., 2008***), adjusting the spatial scale of the warp correction on a session-by-session basis. To mitigate the noise of frame-to-frame warp calculations, *x*- and *y*-warp fields were adaptively filtered in time. Comparisons of denoised and warp corrected images can be seen in Figure 2—figure supplement 2B-D. Due to their relatively larger size and spherical shape, soma imaging did not require warp correction.

#### Segmentation and demixing

Segmentation and demixing were performed on an image series comprising all imaging sessions from a given animal and given layer. This image series was cropped to contain only pixels with valid data across the entire image series (Figure 2—figure supplement 2B). We used a software package optimized for performing NMF on large datasets within a distributed computing environment (***Chennupati et al., 2020***) to calculate a single factorization (Figure 2—figure supplement 2A). NMF was performed on a compressed version (*N* = 3 frame binning), normalized, and baseline-corrected version of the image series (see Figure 2—figure supplement 2A). Unlike piecemeal approaches to NMF-based segmentation of large dendritic imaging datasets (***Buchanan et al., 2018***), the output of this complete NMF pipeline was not prone to spatial discontinuity artifacts from block stitching. We calculated a single factorization of at least 100 initial components within each FOV to ensure that even weakly active dendrites or somata would be identified. Minimally active components and components with a spatial distribution indicating that they were noise-dominated were excluded from final components. Initial components with temporal correlation > 0.8, spatial correlation > 0.5, and binarized spatial overlap > 50% were merged to generate a final set of ROIs (Figure 2G). For each imaging session, temporal components were calculated from denoised and warp-corrected image series using Hierarchical Alternating Least Squares (HALS; python scikit-learn; ***Pedregosa et al., 2011***) with the ROIs as fixed spatial components. Finally, a binarized mask derived from the spatial component of each ROI was used to extract a demixed activity trace for each ROI from the matrix product of each ROI’s spatial and temporal components. ROIs were excluded from further analysis if they corresponded to axons in L1, basal dendrites in L5, or were lost intermittently during imaging sessions.

We assume that individual ROIs rarely included segments of dendrite from more than one neuron. Tuft activity is sparse (***Francioni et al., 2019***) and GCaMP8m kinetics are fast (***Zhang et al., 2023***), therefore, the activity of the tuft dendrites of two different neurons would have to be highly temporally correlated (> 0.8 Pearson correlation) to merge into a single ROI. Furthermore, components extracted from individual imaging sessions had consistent spatial structure (Figure 2—figure supplement 2F), suggesting that the structure of ROIs was robust to measurement noise and spurious within-session correlations. However, we do not assume that dendritic segments belonging to the same neuron always merged into one ROI. In particular, L5 ET neurons with early bifurcating apical dendrites can exhibit highly independent activity across two hemi-trees (***Otor et al., 2022***) and may be segmented into two ROIs by our approach. In summary, different ROIs are putatively the dendrites of different neurons or highly independent dendritic compartments of the same neuron.

#### Deconvolution

An initial Δ*F ∕F_0_* was calculated from each ROI’s demixed activity trace, where *F*_0_ was the 20^th^ percentile of *F* within a rolling 100 s window. Residual contamination of the estimated *F*_0_ by periods of high activity was corrected by iteratively identifying periods of activity (> 1.0 s.d. of the noise) in a high-pass (1 Hz) filtered version of Δ*F ∕F_0_*, and then excluding those periods from the next iteration of *F*_0_ estimation. Corrected Δ*F ∕F* traces were then deconvolved using constrained FOOPSI (CaImAn; ***Giovannucci et al., 2019***) set to fit an AR(2) model. Deconvolved activity below a noise threshold (defined by bimodal Gaussian fitting of all deconvolved values; python scipy, ***Virtanen et al., 2020***) was set to zero. The resulting graded “impulse” traces were used for all subsequent analysis.

For Figure 2—figure supplement 3, noise was estimated as the square-root of the geometric mean of the Welch power spectrum in a high-frequency band (0.25–0.5 times the frame rate; ***Giovannucci et al., 2019***).

### Trial-alignment and ROI grouping

Individual trial traces were aligned to either the GO cue frame (green line/label in all figures) or the first lickport contact frame (FL; dark yellow line/label in all figures). Trial-type averages were calculated using the mean of *N* = 1000 bootstraps with replacement. For Figure 3G,H, ROIs were excluded if they had *N* < 10 pre-shift CR trials, *N* < 10 pre-shift CL trials, or *r* < 0.5 correlation between trial averages of 50/50 trial count splits (***Yang et al., 2022***). In Figure 3G,H, ROIs were classified and sorted according to the timing of peak activity and whether this peak activity was in CR or CL GO-aligned averages. For Figure 5D,F, ROIs were excluded if they had *N* < 10 CA trials and sorted according to mean *CR*−*CA* activity between 0 s and 1 s relative to first-lick contact (see Figure 5—figure supplement 1).

### Identification of GO- and contact-associated activity

The deconvolved activity of each ROI was modeled as the sum of GO- and contact-aligned impulse response functions (IRFs). In brief, vectors marking GO-cue times and first lick tongue contact times (FL) across all pre-shift CR or pre-shift CL trials were converted into lagged Toeplitz design matrices. These design matrices were then fit to the activity of each ROI using non-negative linear regression and the resulting coefficients were taken as the estimated IRFs. To compare the contribution of GO-and contact-associated activity with respect to a common temporal alignment, all contact-IRFs for a given animal were shifted by the temporal offset between the GO-cue and the mean FL time for that trial-type (CR or CL) and animal. Thus, both GO- and contact-associated trial-aligned activity were realigned to the timing of the GO-cue. Mean CR and mean CL activities were then averaged together, generating a single GO-associated activity trace and a single contact-associated activity trace for each ROI. To compare relative peri-GO activity across dendrites and somata (Figure 4B,C), activity was divided by the mean of both GO-associated and contact-associated activity during the first half (0.625 s) of the delay epoch. Population averages of contact-IRFs that were not shifted prior to averaging were nearly identical (excluding the overall temporal shift; data not shown), indicating that pooling of mean IRFs across animals and trial types produces minimal smearing of the final population IRF.

### Coding directions and projections of task-variables

The mean activity of ROIs during each trial type was modeled as a linear combination of three task variables: stimulus, choice, and outcome (***Chen et al., 2017***; ***Yang et al., 2022***). At each timepoint *t* in the trial, three task-variable selectivity CDs were defined as:

- Stimulus CD:

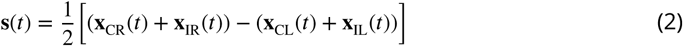
- Choice CD:

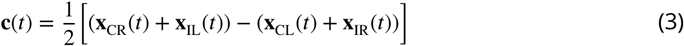
- Outcome CD:

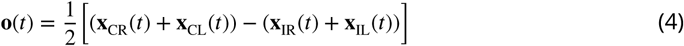

where **x***_k_*(*t*) denotes the trial-averaged activity for trial type *k* at time *t*. For each timepoint, each selectivity CD was then orthogonalized with respect to the other two selectivity CDs by a QR decomposition in which that selectivity CD was last in the order.

We then projected trial-averaged activity at *t* onto the corresponding **s**(*t*), **c**(*t*), and **o**(*t*) CDs. To cross-validate projections, CDs were calculated from a randomly selected 50% of trials and then the remaining 50% of trials were projected onto the CD. Population selectivity along each CD at *t* was then calculated by replacing **x***_k_*(*t*) in Equations 2–4 with the corresponding projections of trial-averaged activity along each CD. For estimation of SEM and statistical comparisons, data were hierarchically bootstrapped at the top level (animals) *N_boot_* = 10000 times and 50/50 trial splits were recalculated each bootstrap iteration.

### Coding directions and projections that discriminate trial types

To compare population dynamics between two trial types, we calculated the CDs along which activity maximally discriminated between the trial types (***Stopfer et al., 2003***; ***Li et al., 2016***). CDs were defined as:

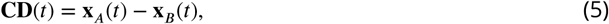

where **x***_A_*(*t*) and **x***_B_*(*t*) denote the trial-averaged activity for trial types *A* and *B* at time *t*.

To calculate selectivity projections, **x***_A_*(*t*) and **x***_B_*(*t*) were projected onto **CD**(*t*) and then subtracted from each other. All projections were cross-validated by calculating CDs from a randomly selected 50% of trials and then projecting onto the remaining 50% of trials. For estimation of SEM and statistical comparisons, data were hierarchically bootstrapped at the top level (animals) *N_boot_* = 10000 times and 50/50 trial splits were recalculated each bootstrap iteration.

#### Coding Magnitude and Correlation

To quantify the overall strength of population selectivity, we computed the *magnitude* of the coding direction at each timepoint *t* as the L2-norm of the CD vector.

To quantify the *correlation* between CDs across training epochs, we calculated the cosine similarity between the mean-subtracted CDs, which was equivalent to the Pearson correlation. Empirical measurements of correlation will be systematically lower than true underlying correlations due to noise. To place across-epoch correlations into context, we estimated a maximum observable correlation (*i.e.*, correlation “ceiling”) based on the within-epoch reliability of the CD estimate. Within-epoch reliability was defined as the correlation between two CDs derived from a random 50/50 trial-level split of the epoch. The maximum observable cross-epoch correlation was defined as:

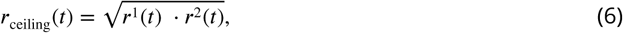

where *r*^1^(*t*) and *r*^2^(*t*) denote the within-epoch correlations of training epoch 1 and training epoch 2. The cross-epoch correlation (*r*_cross_(*t*)) was then normalized by this correlation ceiling to obtain a reliability-corrected correlation estimate:

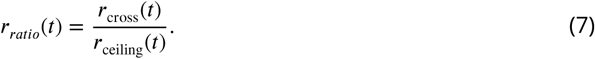

### Selectivity Index, Responsiveness, and Net Selectivity

To quantify the selectivity of individual ROIs (Figure 6C-G), we first calculated the mean trial-averaged activity of each ROI during non-overlapping 0.5 s bins spanning the trial duration. At each time bin *t* we computed a *Selectivity Index* (SI) for ROI and for each training epoch (pre-shift or post-shift):

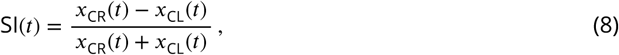

where *x*_CR_(*t*) and *x*_CL_(*t*) are the trial-averaged activities at time bin *t* for CR and CL trials, respectively. The 95% confidence intervals (CIs) for each SI(*t*) were estimated by resampling trials with replacement (*N*_boot_ = 10000). When the number of CR and CL trials differed across training epochs, trials were randomly subsampled to ensure matching trial counts and balanced estimates. SI(*t*) estimates with 95% CI width > 0.66 or minimal total activity (*x*_CR_(*t*) + *x*_CL_(*t*) < 0.1 impulses) were considered unreliable and were excluded. Within a training epoch, ROIs with any reliable SI(*t*) estimates were considered “task-modulated”. For each ROI, the time bin *t* across both training epochs that had the largest difference in activity between the two trial types (*x*_CR_(*t*) − *x*_CL_(*t*)) and a reliable SI(*t*) was defined as the characteristic response time, *t*_response_, for that ROI. The final pre-shift SI and post-shift SI for each ROI were defined as SI(*t*_response_) drawn from the respective SI(*t*) estimates of each epoch.

The characteristic response time (*t*_response_) was then used to calculate the *Responsiveness* and *Net Selectivity* for each ROI:

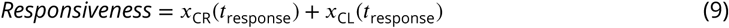

and

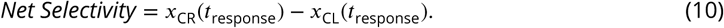

### Sample Sizes

Relevant sample sizes can be found in the appropriate figure legends. Overall sizes for the numbers of animals and ROIs can be found in Table 1 while numbers of sessions and trials can be found in Table 2 and Table 3 respectively.

**Table 1.** Number of animals and ROIs included in each analysis comparing compartments.

| Figure | Dendrites |  | Somata |  |
| --- | --- | --- | --- | --- |
|  | Animals | ROIs | Animals | ROIs |
| Figure 4B-C,G-I | 15 | 234 | 12 | 328 |
| Figure 5H,I | 10 | 167 | 8 | 179 |
| Figure 6C-G | 10 | 169 | 8 | 180 |
| Figure 6I-K | 10 | 167 | 8 | 179 |
| Figure 6—figure supplement 2B-E | 8 | 161 | 9 | 211 |
| Figure 6—figure supplement 3B-E | 8 | 161 | 9 | 211 |

**Table 2.** Number of sessions by training epoch.

| Training Epoch | Median Sessions | Standard Deviation |
| --- | --- | --- |
| Pre-shift early | 2.0 | 0.29 |
| Pre-shift late | 2.0 | 0.00 |
| Post-shift early | 2.0 | 0.22 |
| Post-shift late | 2.0 | 0.42 |

**Table 3.**
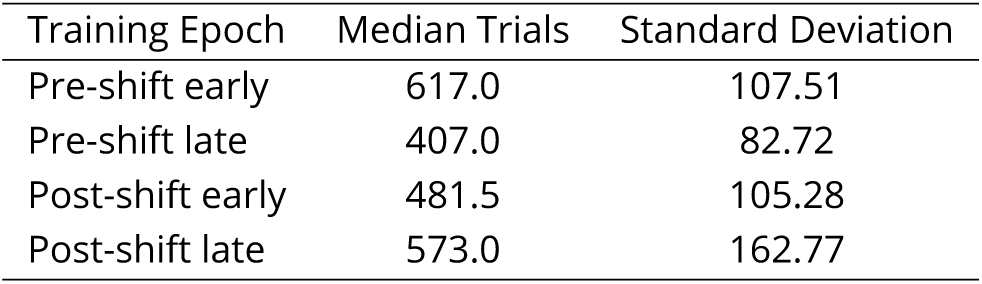
Number of trials by training epoch.

| Training Epoch | Median Trials | Standard Deviation |
| --- | --- | --- |
| Pre-shift early | 617.0 | 107.51 |
| Pre-shift late | 407.0 | 82.72 |
| Post-shift early | 481.5 | 105.28 |
| Post-shift late | 573.0 | 162.77 |

### Binning and Statistics

In all plots, the dendrite and soma data were binned with matching duration time bins. Data were binned at: 44 ms (Figure 3; Figure 4D,E; Figure 5C,E), 90 ms (Figure 4B), 224 ms (Figure 4F; Figure 5D,F,H,I; Figure 5—figure supplement 1; Figure 5—figure supplement 2; Figure 6I,J; Figure 6—figure supplement 1B; Figure 6—figure supplement 2B; Figure 6—figure supplement 3B) or 493 ms (Figure 6C-G). Unless otherwise noted, data are presented as bootstrap mean ± standard error of *N*_boot_ = 1000 bootstraps for trace averages and *N*_boot_ = 10000 bootstraps for statistical tests. Unless otherwise noted, * indicates *p* < 0.05 estimated by hierarchical bootstrap at the level of animals. Horizontal gray dashed lines indicate the comparison value for significance tests. Corrections for multiple comparisons were performed using a Holm–Bonferroni method. Comparisons between probability distributions were performed using a Kolmogorov-Smirnov (K-S) test (Figure 2—figure supplement 3B,C).

## Code and data availability

Code and data intermediates used in the generation of this manuscript are available at https://github.com/kerlin-lab/Scheib_2026. The complete raw image series are ≈40 TB and will be available on request.

## Acknowledgments

We thank Timothy Ebner, Matthew Chafee, and Thomas Naselaris for comments on the manuscript. This work was supported with funding from the Whitehall Foundation (Grant 2020-05-71), NINDS R01NS127902, as well as the resources and staff at the University Imaging Centers (UIC; RRID: SCR_020997) and Minnesota Supercomputing Institute (MSI) at the University of Minnesota.

---

**Figure 1—figure supplement 1.**
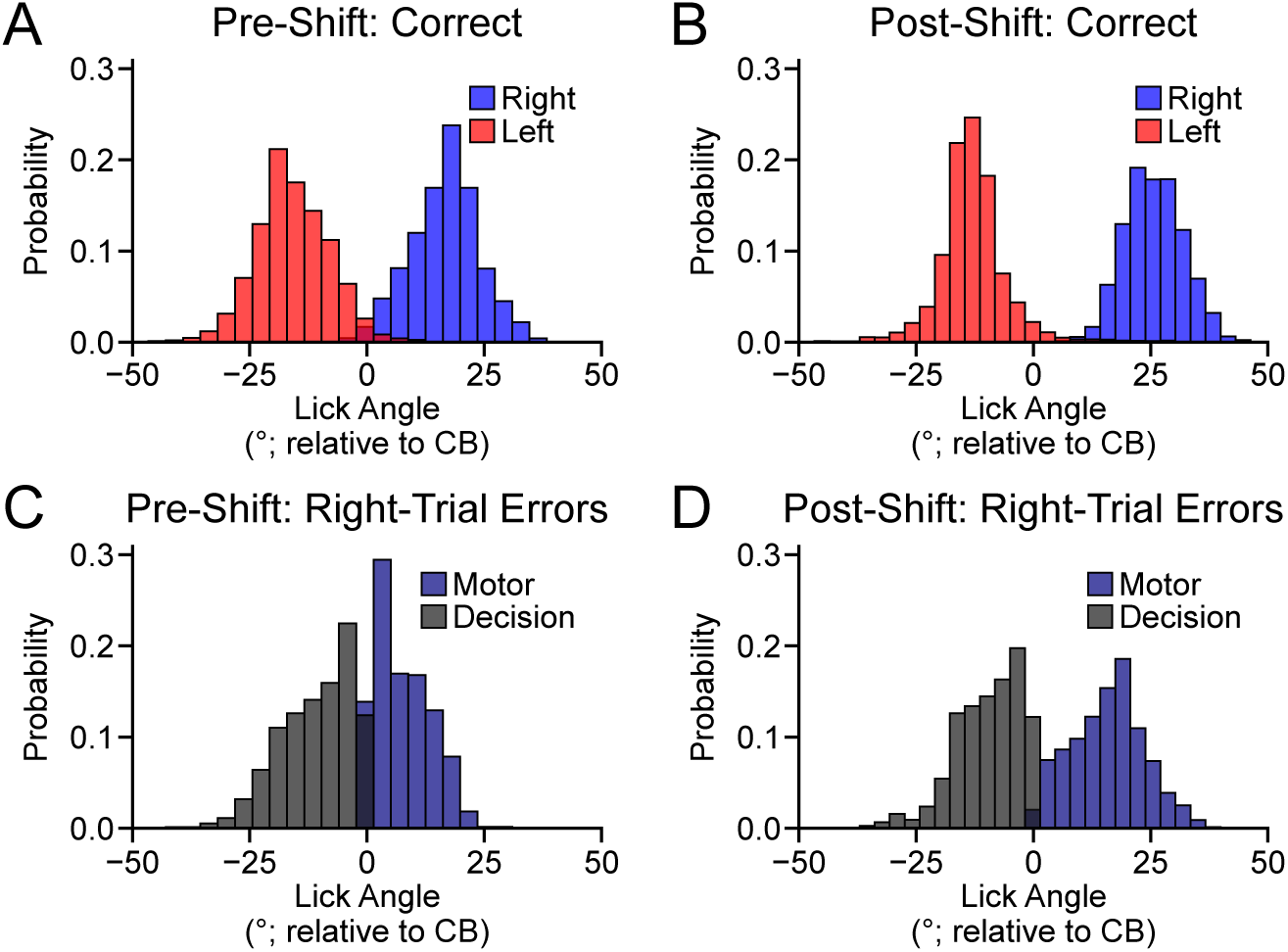
Tongue exit angles change after the port shift to avoid the obstructing lickport. (A) Empirical probability distributions for pre-shift correct trial types across all animals (pre-shift Correct Right: 16.6°± 0.1, *N* = 10610; pre-shift Correct Left: −15.8°± 0.1, *N* = 9302). (B) Distributions of post-shift correct trials (post-shift Correct Right: 25.6°± 0.1, *N* = 6568; post-shift Correct Left: −12.9°± 0.1, *N* = 4717). (C) Distributions of pre-shift right error trials (pre-shift Motor Error Right: 7.7°± 0.2, *N* = 649; pre-shift Decision Error Right: −10.8°± 0.2, *N* = 1534). The distribution of all pre-shift error exit angles was unimodal (ΔBIC = −28.3, Gaussian mixture model comparison, see Methods for details). (D) Distributions of post-shift right error trials (post-shift Motor Error Right: 15.8°± 0.1, *N* = 4010; post-shift Decision Error Right: −9.6°± 0.3, *N* = 754). The distribution of post-shift error exit angles was bimodal (ΔBIC = 675.5). All statistics are mean ± SEM.

**Figure 2—figure supplement 1.**
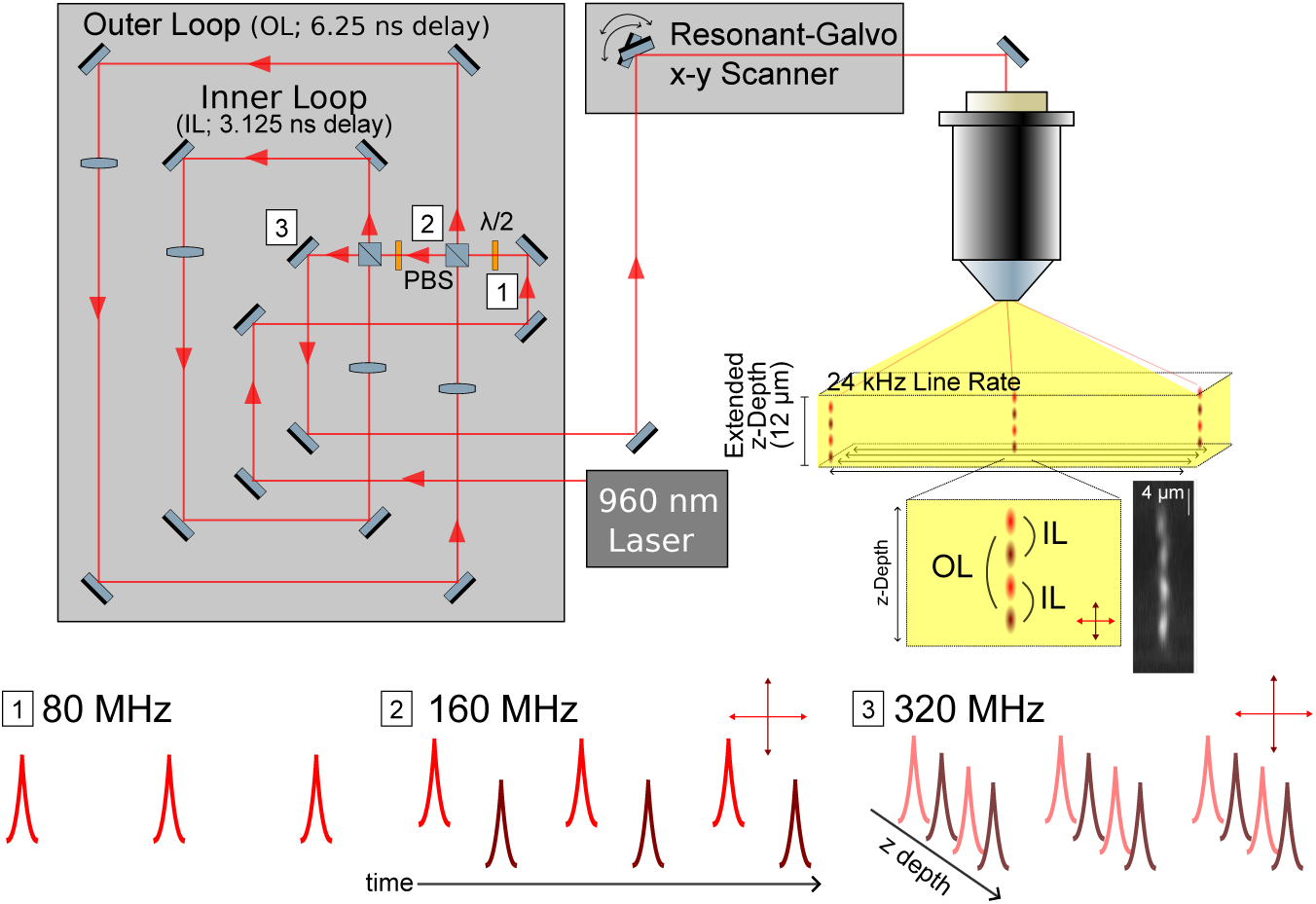
Optical design of module for extended axial 2P imaging. A linearly polarized 80 MHz laser pulse train (1) was split into polarization-dependent paths *via* a polarizing beam splitter (PBS), where a half-wave plate (λ∕2) controls the distribution of energy to each path. The *S*-polarized component incurred a 6.25 ns delay along the *Outer Loop* with respect to the un-delayed *P*-polarized component, which was then recombined using the same PBS to form a 160 MHz pulse train with alternating polarization (2). Next, the global polarization of the 160 MHz pulse train was rotated such that the *S*-polarized components of each orthogonally polarized element of the 160 MHz pulse train incurred a delay of 3.125 ns in propagation along an *Inner Loop* with respect to the *P*-polarized components. Pulses were then recombined *via* a PBS resulting in a 320 MHz pulse train (3). Additionally, the *Outer Loop* and *Inner Loop* both contained 1:1 optical relays arranged to impart a variable divergence such that each unique path through the pulse splitter generated a distinct downstream shift in focal plane. Each of these paths were conjugated to the back aperture of the objective lens through a series of optical relays. A home-built single prism pulse compressor was used to compensate the average group delay dispersion of all delay paths and the rest of the microscope.

**Figure 2—figure supplement 2.**
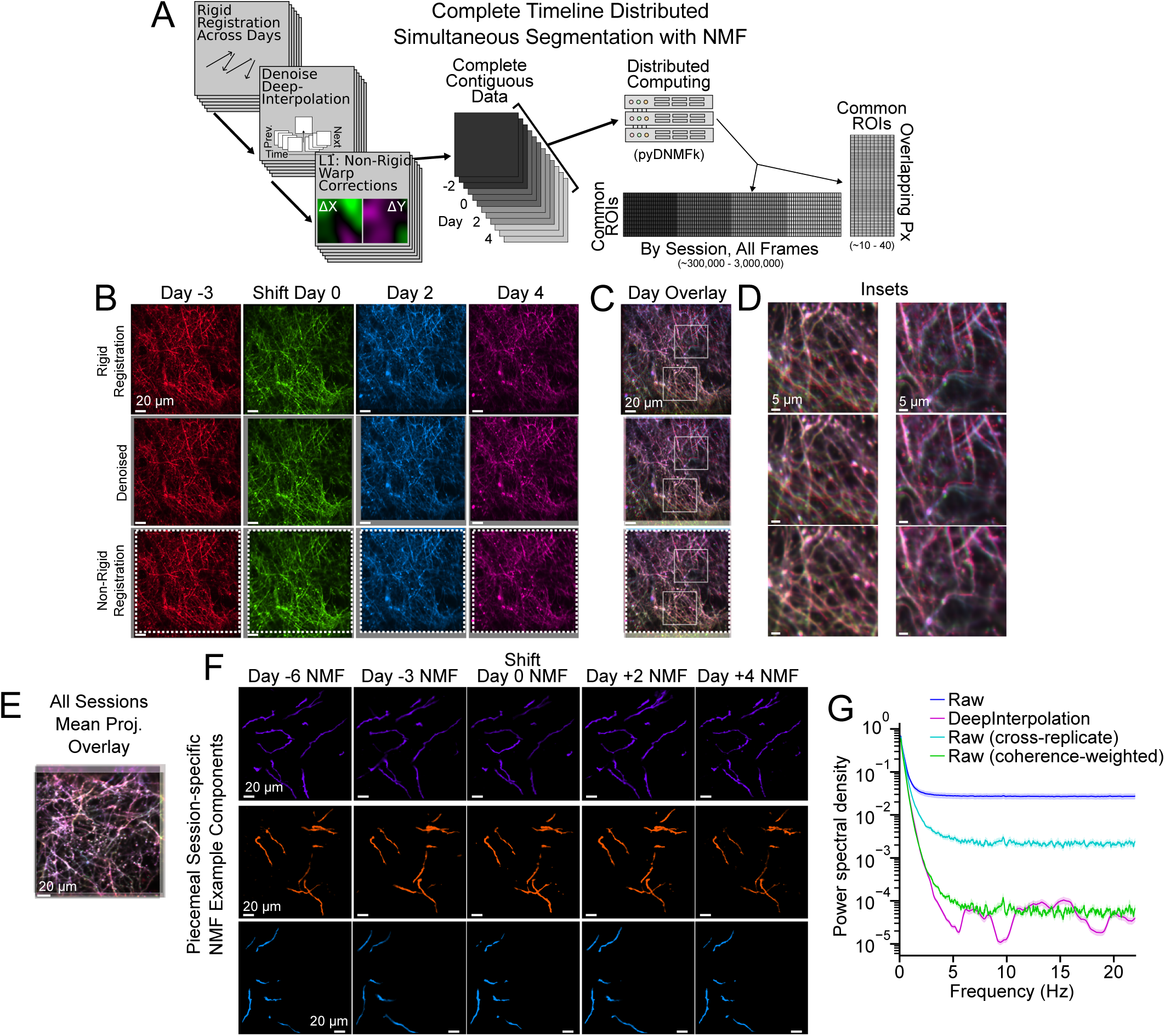
Image registration, denoising, and segmentation pipeline. (A) Schematic representation of the image analysis pipeline including rigid registration, denoising, non-rigid warp correction, and then complete timeline NMF. (B) Mean projection images for four dendrite imaging sessions showing the rigid registration output (top row), denoised output (middle row) and the non-rigid registration output (bottom row). Gray border areas are regions where the data were cropped prior to denoising to eliminate any regions that did not have consistent data throughout the imaging session following the rigid registration. White dotted box indicates the maximum shared region where no data was missing due to registration across all four imaging sessions. (C) Overlay of all four sessions for the corresponding rigid (top), denoised (middle) and non-rigid (bottom) corrected mean projections in (B). (D) Insets for regions in (C) showing that after rigid registration and denoising there are sometimes offsets in the positions of individual dendritic branches that must be corrected with the non-rigid registration. (E) Mean projection image overlay across all recorded sessions for an example dendritic FOV in Figure 2G. (F) Stability of NMF spatial components. Spatial components corresponding to components in Figure 2G, but calculated independently for each behavioral session. (G) Mean power spectral density (PSD) of dendrite timecourses with or without various forms of noise removal or compensation (mean ±SEM; *N* = 161 ROIs, 3 imaging sessions, 3 animals). Raw: PSD of timecourses extracted from motion-corrected image series. DeepInterpolation: same as Raw, but with denoising. Raw (cross-replicate): estimated “latent” PSD calculated as the mean cross spectral density (CSD) of random 50:50 splits of each dendrite’s ROI pixels (*i.e.,* replicates). Raw (coherence-weighted): estimated “recoverable” PSD calculated as the mean PSD of the raw timecourses weighted by the coherence of replicates.

**Figure 2—figure supplement 3.**
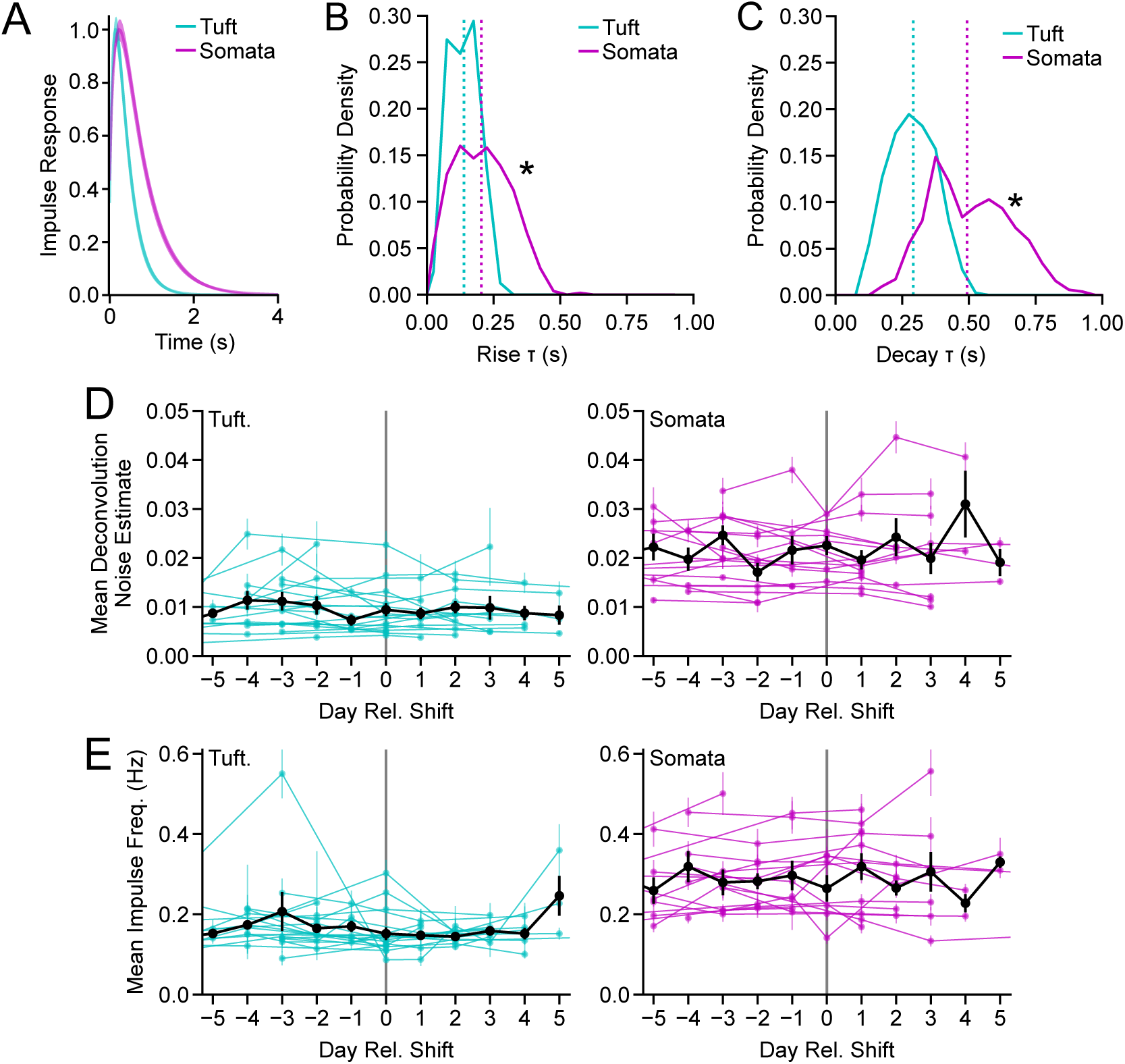
Estimated event kinetics, noise, and impulse rates across longitudinal 2P imaging. (A) Mean impulse response function (IRF, normalized to peak before averaging) estimated for tuft dendrites (*N* = 402 dendrites from *N* = 21 animals) and somata (*N* = 528 ROIs from *N* = 20 animals) from fits to an AR(2) model using constrained FOOPSI (***Pnevmatikakis et al., 2016***). (B) Probability densities for the rise time constant (*τ*) of the fits. Vertical dotted lines indicate the population median. * indicates *p* < 0.05 Kolmogorov-Smirnov (K-S) test. (C) Same as (B), but for the decay time constant (*τ*). (D) Standard deviation of the noise averaged across ROIs of each animal by imaging session for 5 days prior to port shift and 5 days following port shift in dendrites (left) and somata (right). Black line indicates the average noise estimate across all animals. Noise was estimated from high frequencies in the power spectral density of the trace of each ROI (***Pnevmatikakis et al., 2016***). (E) Same as (D) but for estimated impulse rates after thresholding the deconvolved traces (see Methods). Error bars: ±SEM (not bootstrapped).

**Figure 4—figure supplement 1.**
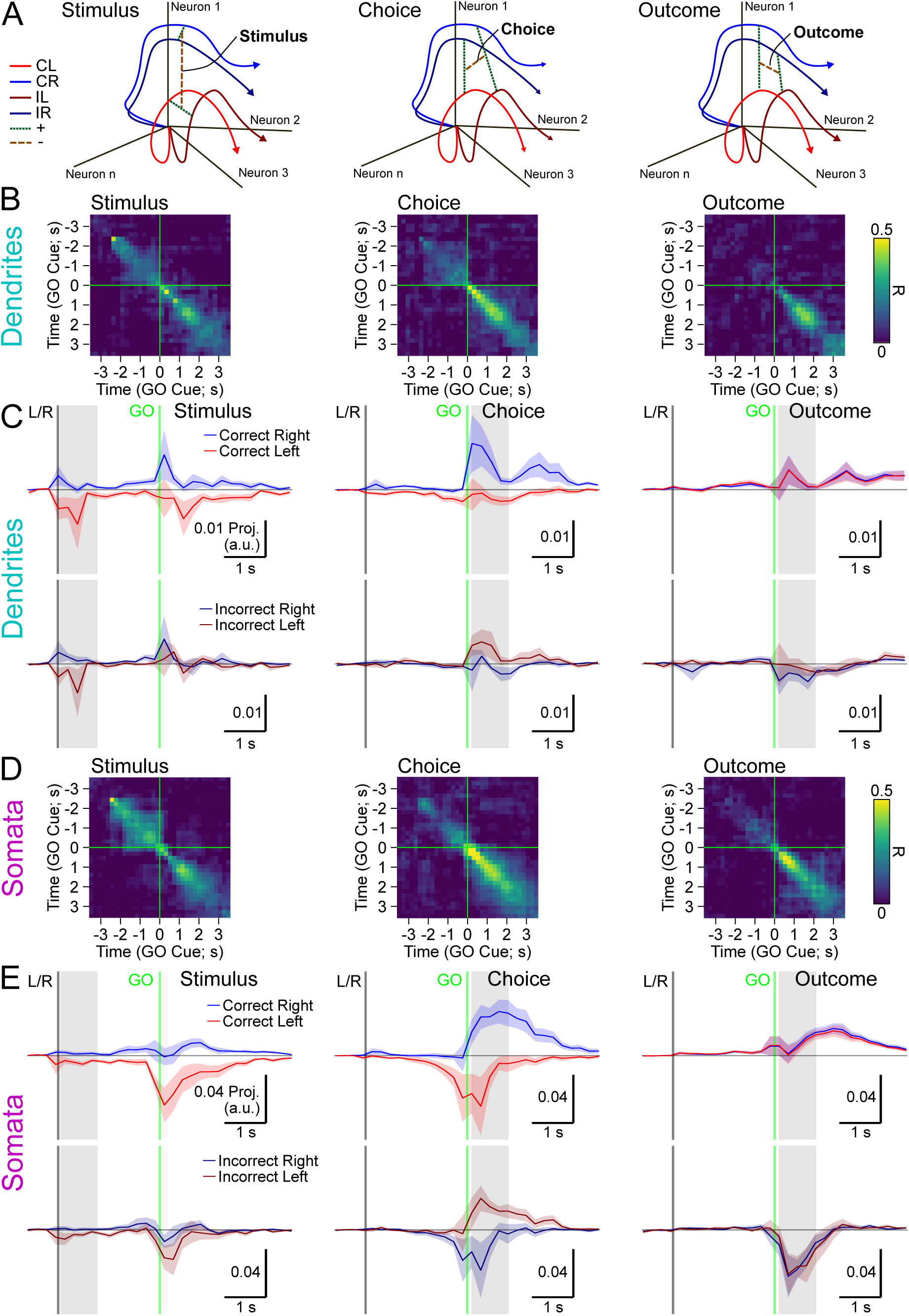
Mean trial-type projections along *Stimulus*, *Choice*, and *Outcome* CDs. (A) Illustration of the calculation of *Stimulus*, *Choice*, and *Outcome* CDs within activity space. For each CD, dotted lines indicate trial types that were first summed and the dash-dotted line indicates subsequent subtraction of the results to obtain the CD. (B) Cross-validated (50:50 trial split) correlation of CD weights across *Stimulus*, *Choice*, and *Outcome* dimensions for dendrites. (C) Cross-validated projections of trial-averaged activity. Correct and incorrect trial projections shown in the top and bottom of each panel, respectively. (D) Same as (B), but for all somata. (E) Same as (C), but for all somata. Error bars: hierarchical bootstrap SEM.

**Figure 5—figure supplement 1.**
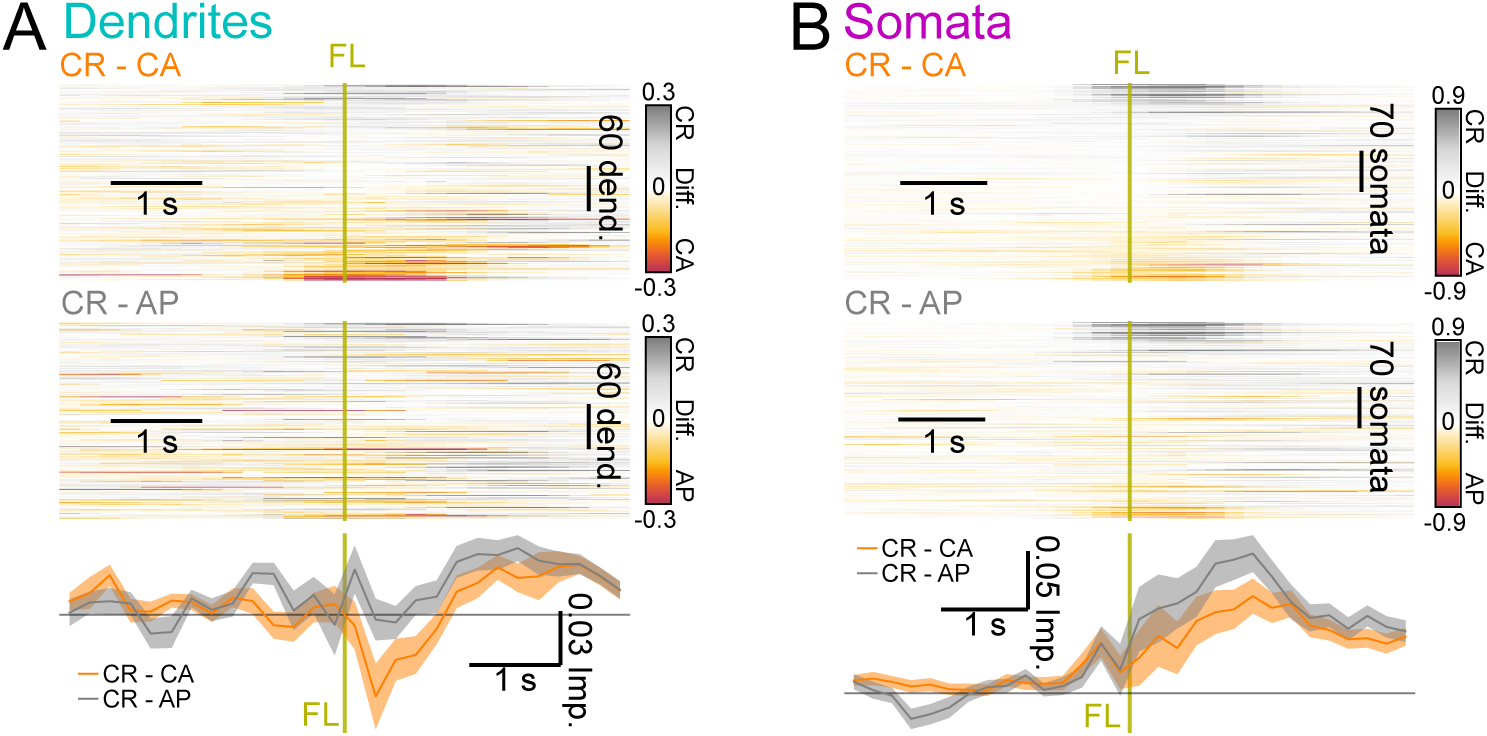
Differences in mean trial-aligned CR, CA, and AP activity. (A) All individual ROI differences in the mean CR minus the mean CA activity (top) or mean CR activity minus mean AP activity (bottom). ROI order is same as in Figure 5D. (B) Same as (A) for somata from Figure 5F.

**Figure 5—figure supplement 2.**
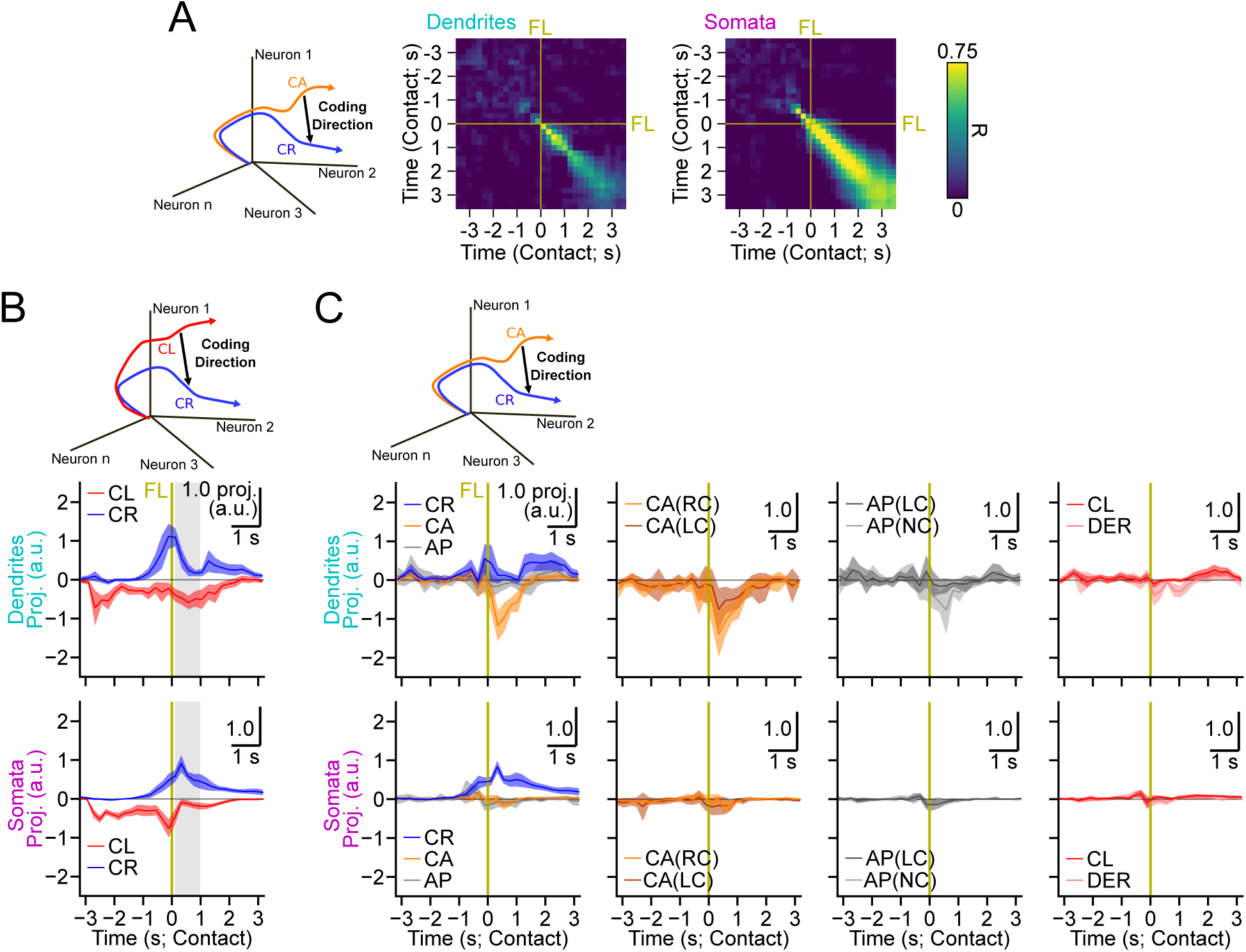
*CR* − *CA* code stability and additional trial-type projections. (A) Illustration of *CR* − *CA* CD calculation (left) and cross-validated correlations of *CR* − *CA* CD across trial time relative to first lickport contact (FL; dark yellow line). (B) Illustration of *CR* −*CL* CD calculation (top), as well as dendrite (middle) and somata (bottom) normalized cross-validated projections along the *CR* − *CL* CD. Projections were normalized by division by *CR* + *CL* in shaded region. (C) Illustration of *CR* − *CA* CD calculation (top), as well as dendrite (middle) and somata (bottom) normalized cross-validated projections along the *CR* − *CA* CD for additional trial-types. Projections were normalized by division by *CR* + *CL* in shaded region of (B) to allow for direct comparisons of magnitude. Trial-type abbreviations are CR: Correct Right; CL: Correct Left; CA: Correction Attempted; AP: Abandoned Port; CA(RC): Correction Attempted (second lick made Right port Contact); CA(LC): Correction Attempted (second lick made Left port Contact); AP(LC): Abandoned Port (second lick made Left port Contact); AP(NC): Abandoned Port (second lick did Not make Port contact); DER: Decision Error Right.

**Figure 6—figure supplement 1.**
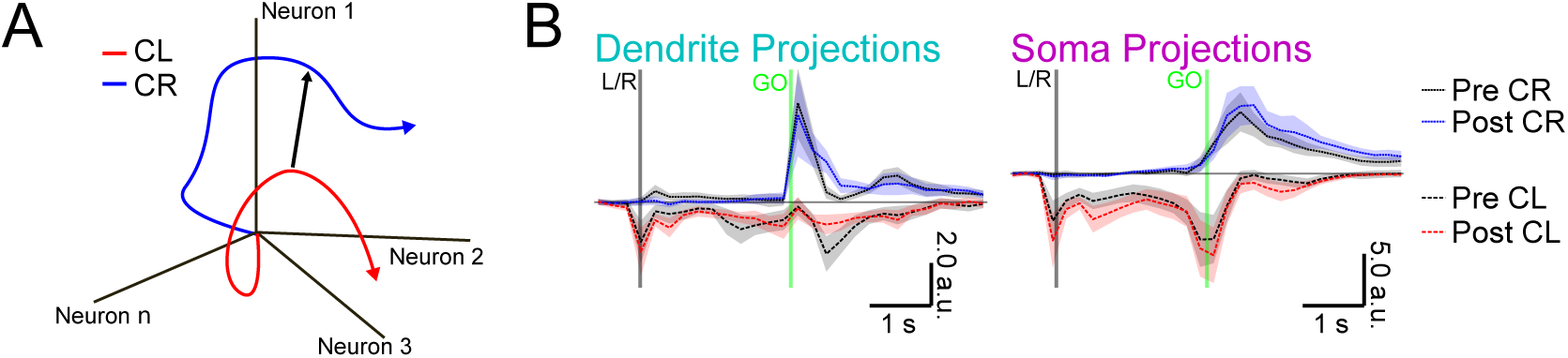
Trial-type projections before and after the lickport shift. (A) Schematic of the *CR*−*CL* CD. (B) Cross-validated projections of CR and CL trials along the *CR*−*CL* CD. Same data as used for Figure 6I,J. Shaded error: hierarchical bootstrap SEM.

**Figure 6—figure supplement 2.**
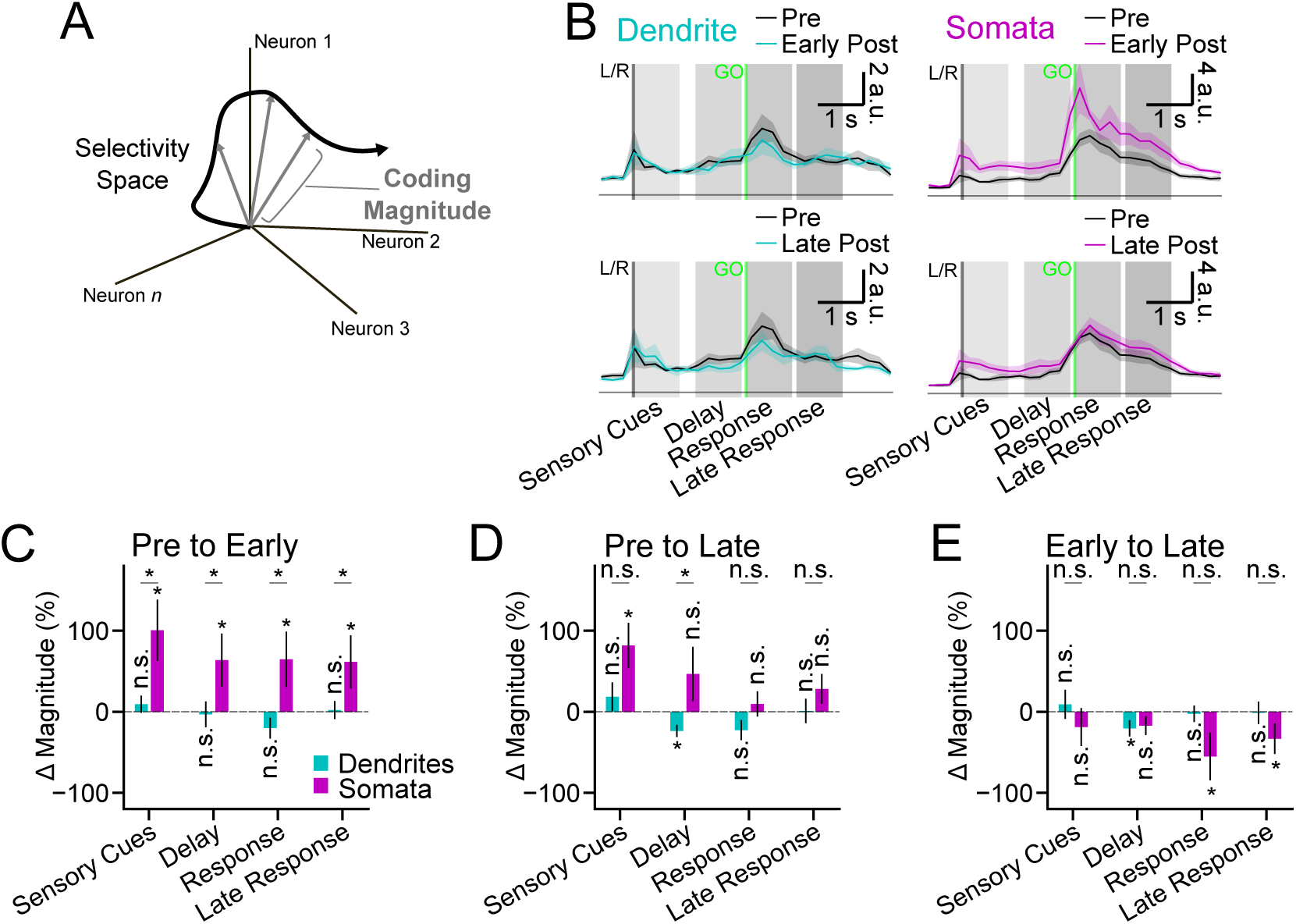
Selectivity CD magnitude across early and late learning. (A) Illustration of the calculation of the coding magnitude in selectivity space. (B) Coding magnitude of dendrites (left) and somata (right) during pre-shift (black), early post-shift (top row), and late post-shift (bottom row) training epochs. (C) Percent change in coding magnitude from pre-shift to early post-shift averaged within the four time windows indicated by gray rectangles in (B). (D) Same as (C), but from pre-shift to late post-shift. (E) Same as (C) but from early post-shift to late post-shift. Shaded error and error bars: hierarchical bootstrap SEM.

**Figure 6—figure supplement 3.**
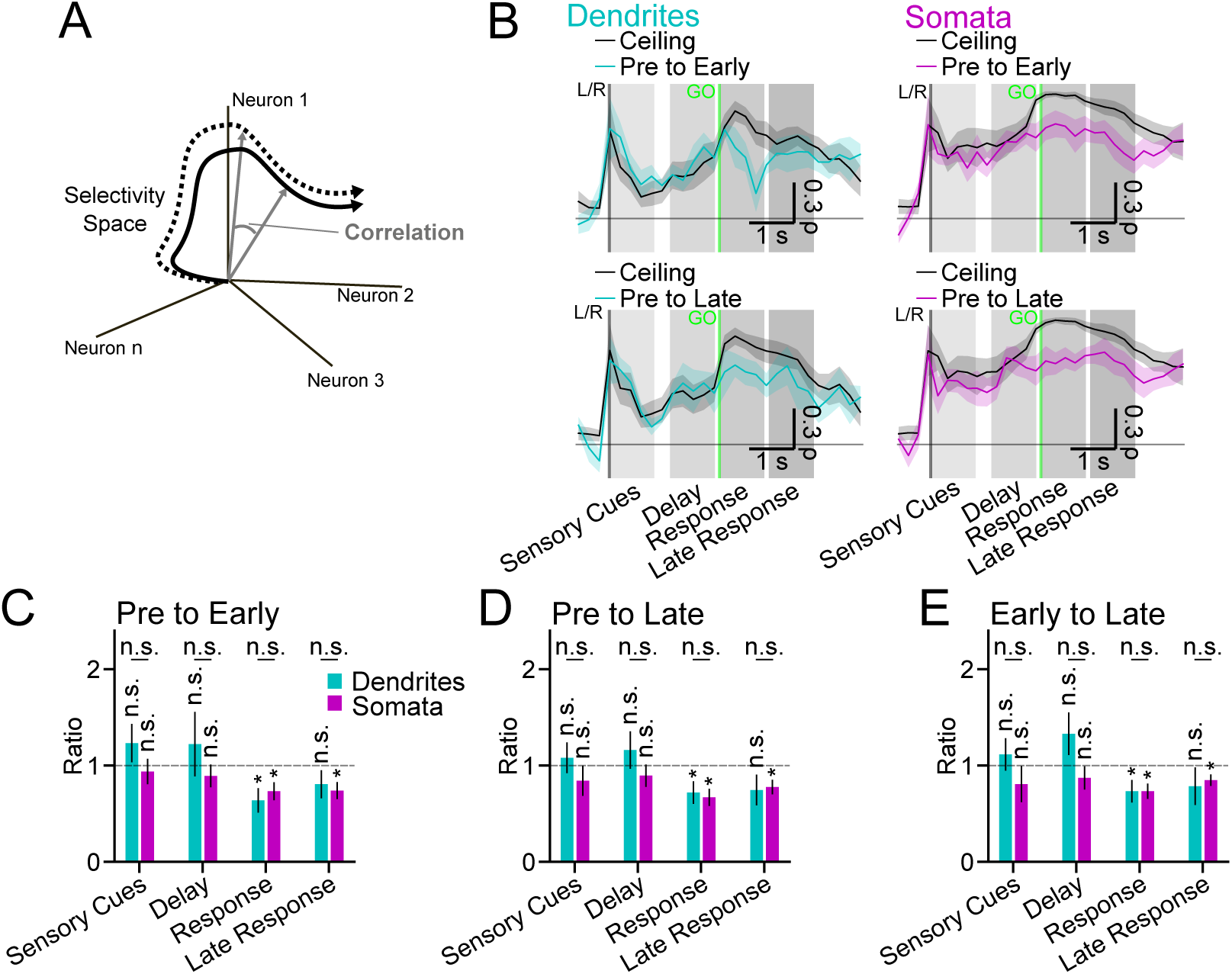
Selectivity CD correlations across early and late learning. (A) Illustration of correlation (cosine similarity) calculation between CDs in selectivity space. (B) Correlation between CDs calculated from pre-shift activity and CDs calculated from early post-shift activity for dendrites (top left, cyan line) and somata (top right, magenta line), as well as correlation between CDs calculated from pre-shift activity and CDs calculated from late post-shift activity for dendrites (bottom left, cyan line) and somata (bottom right, magenta line). Black lines indicate the estimated maximum possible correlation (“ceiling”) across the training epochs given correlations of repeated measures within each epoch (*i.e.*, the limit due to degradation by measurement noise; see Methods). (C) Ratios between the early post-shift to pre-shift correlations and the correlation ceilings in (B) averaged within the four time windows indicated by gray rectangles in (B). A ratio ≈1 indicates no change in the CD pattern across the training epochs beyond measurement noise, whereas a ratio significantly < 1 indicates significant change across the training epochs in the CD pattern. (D) Same as (C), but for correlation of late post-shift to pre-shift. (E) Same as (C), but for correlation of late post-shift to early post-shift. Shaded error and error bars: hierarchical bootstrap SEM.

## References

1. Ako R, Terada SI, Matsuzaki M. A cerebello-thalamo-cortical pathway transmits reward-based post-error signals for motor timing correction during learning in male mice. Nature Communications. 2025 Aug; 16(1):7663. https://www.nature.com/articles/s41467-025-62831-6, doi: 10.1038/s41467-025-62831-6.

2. Ames KC, Ryu SI, Shenoy KV. Simultaneous motor preparation and execution in a last-moment reach correction task. Nature Communications. 2019 Jun; 10(1):2718. https://www.nature.com/articles/s41467-019-10772-2, doi: 10.1038/s41467-019-10772-2.

3. Avants BB, Epstein CL, Grossman M, Gee JC. Symmetric diffeomorphic image registration with cross-correlation: Evaluating automated labeling of elderly and neurodegenerative brain. Medical Image Analysis. 2008 Feb; 12(1):26–41. https://www.sciencedirect.com/science/article/pii/S1361841507000606, doi: 10.1016/j.media.2007.06.004.

4. Benezra SE, Patel KB, Perez Campos C, Hillman EM, Bruno RM. Learning enhances behaviorally relevant representations in apical dendrites. eLife. 2024 Dec; 13:RP98349. 10.7554/eLife.98349, doi: 10.7554/eLife.98349.

5. Bollu T, Ito BS, Whitehead SC, Kardon B, Redd J, Liu MH, Goldberg JH. Cortex-dependent corrections as the tongue reaches for and misses targets. Nature. 2021 Jun; 594(7861):82–87. https://www.nature.com/articles/s41586-021-03561-9, doi: 10.1038/s41586-021-03561-9.

6. Buchanan K, Kinsella I, Zhou D, Zhu R, Zhou P, Gerhard F, Ferrante J, Ma Y, Kim S, Shaik M, Liang Y, Lu R, Reimer J, Fahey P, Muhammad T, Dempsey G, Hillman E, Ji N, Tolias A, Paninski L. Penalized matrix decomposition for denoising, compression, and improved demixing of functional imaging data. bioRxiv. 2018; 10.1101/334706, doi: 10.1101/334706.

7. Maristany de las Casas E, Killmann K, Drüke M, Münster L, Ebner C, Sachdev R, Jaeger D, Larkum ME. Tuft dendrites in frontal motor cortex enable flexible learning. Science. 2026 May; 392(6798):eadx4358. https://www.science.org/doi/10.1126/science.adx4358, doi: 10.1126/science.adx4358, publisher: American Association for the Advancement of Science.

8. Chen TW, Li N, Daie K, Svoboda K. A Map of Anticipatory Activity in Mouse Motor Cortex. Neuron. 2017 May; 94(4):866–879.e4. https://www.cell.com/neuron/abstract/S0896-6273(17)30406-3, doi: 10.1016/j.neuron.2017.05.005.

9. Chennupati G, Vangara R, Skau E, Djidjev H, Alexandrov B. Distributed non-negative matrix factorization with determination of the number of latent features. The Journal of Supercomputing. 2020 Sep; 76(9):7458–7488. 10.1007/s11227-020-03181-6, doi: 10.1007/s11227-020-03181-6.

10. Churchland MM, Yu BM, Ryu SI, Santhanam G, Shenoy KV. Neural Variability in Premotor Cortex Provides a Signature of Motor Preparation. Journal of Neuroscience. 2006 Apr; 26(14):3697–3712. https://www.jneurosci.org/content/26/14/3697, doi: 10.1523/JNEUROSCI.3762-05.2006.

11. Cichon J, Gan WB. Branch-specific dendritic Ca2+ spikes cause persistent synaptic plasticity. Nature. 2015 Apr; 520(7546):180–185. https://www.nature.com/articles/nature14251, doi: 10.1038/nature14251.

12. Costa RM, Cohen D, Nicolelis MAL. Differential Corticostriatal Plasticity during Fast and Slow Motor Skill Learning in Mice. Current Biology. 2004 Jul; 14(13):1124–1134. https://linkinghub.elsevier.com/retrieve/pii/S0960982204004658, doi: 10.1016/j.cub.2004.06.053.

13. Daie K, Wang T, Singh A, Finkelstein A, Kim JJ, Rozsa M, Svoboda K. ALM Window Surgery. protocolsio. 2023; https://www.protocols.io/view/alm-window-surgery-4r3l24qojg1y/v1, doi: 10.17504/protocols.io.bqstmwen.

14. Dasgupta R, Dong M, O’Connor DH. Flexible sensitivity to inputs during skilled tongue movements. bioRxiv. 2025 May; doi: 10.1101/2025.05.18.654702.

15. Dinh T, kerlin-lab/tongue-classification; 2026. https://github.com/kerlin-lab/tongue-classification.

16. Dragoi T, Loschinskey ZF, Hasnain MA, DePasquale B, Economo MN. Dynamic engagement of the motor cortex in controlling movement. bioRxiv. 2026 Jan; doi: 10.64898/2026.01.13.699314.

17. Evarts EV. Relation of pyramidal tract activity to force exerted during voluntary movement. Journal of Neurophysiology. 1968 Jan; 31(1):14–27. doi: 10.1152/jn.1968.31.1.14.

18. Farinella M, Ruedt DT, Gleeson P, Lanore F, Silver RA. Glutamate-Bound NMDARs Arising from In Vivo-like Network Activity Extend Spatio-temporal Integration in a L5 Cortical Pyramidal Cell Model. PLOS Computational Biology. 2014 Apr; 10(4):e1003590. https://journals.plos.org/ploscompbiol/article?id=10.1371/journal.pcbi.1003590, doi: 10.1371/journal.pcbi.1003590.

19. Fişek M, Herrmann D, Egea-Weiss A, Cloves M, Bauer L, Lee TY, Russell LE, Häusser M. Cortico-cortical feedback engages active dendrites in visual cortex. Nature. 2023 May; 617(7962):769–776. https://www.nature.com/articles/s41586-023-06007-6, doi: 10.1038/s41586-023-06007-6.

20. Francioni V, Padamsey Z, Rochefort NL. High and asymmetric somato-dendritic coupling of V1 layer 5 neurons independent of visual stimulation and locomotion. eLife. 2019 Dec; 8:e49145. 10.7554/eLife.49145, doi: 10.7554/eLife.49145.

21. Francioni V, Tang VD, Toloza EHS, Ding Z, Brown NJ, Harnett MT. Vectorized instructive signals in cortical dendrites. Nature. 2026 Feb; p. 1–10. https://www.nature.com/articles/s41586-026-10190-7, doi: 10.1038/s41586-026-10190-7.

22. Garyfallidis E, Brett M, Amirbekian B, Rokem A, Van Der Walt S, Descoteaux M, Nimmo-Smith I. Dipy, a library for the analysis of diffusion MRI data. Frontiers in Neuroinformatics. 2014 Feb; 8. https://www.frontiersin.org/journals/neuroinformatics/articles/10.3389/fninf.2014.00008/full, doi: 10.3389/fninf.2014.00008.

23. Geng HY, Arbuthnott G, Yung WH, Ke Y. Long-range monosynaptic inputs targeting apical and basal dendrites of primary motor cortex deep output neurons. Cerebral Cortex. 2022 Sep; 32(18):3975–3989. doi: 10.1093/cercor/bhab460.

24. Gillon CJ, Pina JE, Lecoq JA, Ahmed R, Billeh YN, Caldejon S, Groblewski P, Henley TM, Kato I, Lee E, Luviano J, Mace K, Nayan C, Nguyen TV, North K, Perkins J, Seid S, Valley MT, Williford A, Bengio Y, et al. Responses to Pattern-Violating Visual Stimuli Evolve Differently Over Days in Somata and Distal Apical Dendrites. The Journal of Neuroscience. 2024 Jan; 44(5):e1009232023. https://www.jneurosci.org/lookup/doi/10.1523/JNEUROSCI.1009-23.2023, doi: 10.1523/JNEUROSCI.1009-23.2023.

25. Giovannucci A, Friedrich J, Gunn P, Kalfon J, Brown BL, Koay SA, Taxidis J, Najafi F, Gauthier JL, Zhou P, Khakh BS, Tank DW, Chklovskii DB, Pnevmatikakis EA. CaImAn an open source tool for scalable calcium imaging data analysis. eLife. 2019 Jan; 8:e38173. 10.7554/eLife.38173, doi: 10.7554/eLife.38173.

26. Guerguiev J, Lillicrap TP, Richards BA. Towards deep learning with segregated dendrites. eLife. 2017 Dec; 6:e22901. 10.7554/eLife.22901, doi: 10.7554/eLife.22901.

27. Guo K, Yamawaki N, Svoboda K, Shepherd GMG. Anterolateral Motor Cortex Connects with a Medial Subdivision of Ventromedial Thalamus through Cell Type-Specific Circuits, Forming an Excitatory Thalamo-Cortico-Thalamic Loop via Layer 1 Apical Tuft Dendrites of Layer 5B Pyramidal Tract Type Neurons. Journal of Neuroscience. 2018 Oct; 38(41):8787–8797. https://www.jneurosci.org/content/38/41/8787, doi: 10.1523/JNEUROSCI.1333-18.2018.

28. Guo ZV, Hires SA, Li N, O’Connor DH, Komiyama T, Ophir E, Huber D, Bonardi C, Morandell K, Gutnisky D, Peron S, Xu Nl, Cox J, Svoboda K. Procedures for Behavioral Experiments in Head-Fixed Mice. PLOS ONE. 2014 Feb; 9(2):e88678. https://journals.plos.org/plosone/article?id=10.1371/journal.pone.0088678, doi: 10.1371/journal.pone.0088678.

29. Guo ZV, Li N, Huber D, Ophir E, Gutnisky D, Ting JT, Feng G, Svoboda K. Flow of Cortical Activity Underlying a Tactile Decision in Mice. Neuron. 2014 Jan; 81(1):179–194. https://www.cell.com/neuron/abstract/S0896-6273(13)00924-0, doi: 10.1016/j.neuron.2013.10.020.

30. Guzulaitis R, Palmer LM. A thalamocortical pathway controlling impulsive behavior. Trends in Neurosciences. 2023 Dec; 46(12):1018–1024. https://www.sciencedirect.com/science/article/pii/S0166223623002199, doi: 10.1016/j.tins.2023.09.001.

31. Heindorf M, Arber S, Keller GB. Mouse Motor Cortex Coordinates the Behavioral Response to Unpredicted Sensory Feedback. Neuron. 2018 Sep; 99(5):1040–1054.e5. https://linkinghub.elsevier.com/retrieve/pii/S0896627318306445, doi: 10.1016/j.neuron.2018.07.046.

32. Hill DN, Varga Z, Jia H, Sakmann B, Konnerth A. Multibranch activity in basal and tuft dendrites during firing of layer 5 cortical neurons in vivo. Proceedings of the National Academy of Sciences. 2013 Aug; 110(33):13618–13623. https://www.pnas.org/doi/abs/10.1073/pnas.1312599110, doi: 10.1073/pnas.1312599110.

33. Hwang EJ, Dahlen JE, Hu YY, Aguilar K, Yu B, Mukundan M, Mitani A, Komiyama T. Disengagement of motor cortex from movement control during long-term learning. Science Advances. 2019 Oct; 5(10):eaay0001. https://www.science.org/doi/10.1126/sciadv.aay0001, doi: 10.1126/sciadv.aay0001.

34. Hwang EJ, Dahlen JE, Mukundan M, Komiyama T. Disengagement of Motor Cortex during Long-Term Learning Tracks the Performance Level of Learned Movements. The Journal of Neuroscience. 2021 Aug; 41(33):7029–7047. https://www.jneurosci.org/lookup/doi/10.1523/JNEUROSCI.3049-20.2021, doi: 10.1523/JNEUROSCI.3049-20.2021.

35. Inagaki HK, Chen S, Daie K, Finkelstein A, Fontolan L, Romani S, Svoboda K. Neural Algorithms and Circuits for Motor Planning. Annual Review of Neuroscience. 2022 Jul; 45(1):249–271. https://www.annualreviews.org/doi/10.1146/annurev-neuro-092021-121730, doi: 10.1146/annurev-neuro-092021-121730.

36. Inagaki HK, Chen S, Ridder MC, Sah P, Li N, Yang Z, Hasanbegovic H, Gao Z, Gerfen CR, Svoboda K. A midbrain-thalamus-cortex circuit reorganizes cortical dynamics to initiate movement. Cell. 2022 Mar; 185(6):1065–1081.e23. https://www.sciencedirect.com/science/article/pii/S0092867422001465, doi: 10.1016/j.cell.2022.02.006.

37. Inagaki HK, Inagaki M, Romani S, Svoboda K. Low-Dimensional and Monotonic Preparatory Activity in Mouse Anterior Lateral Motor Cortex. Journal of Neuroscience. 2018 Apr; 38(17):4163–4185. https://www.jneurosci.org/content/38/17/4163, doi: 10.1523/JNEUROSCI.3152-17.2018.

38. Jiang S, Guan Y, Chen S, Jia X, Ni H, Zhang Y, Han Y, Peng X, Zhou C, Li A, Luo Q, Gong H. Anatomically revealed morphological patterns of pyramidal neurons in layer 5 of the motor cortex. Scientific Reports. 2020 May; 10(1):7916. https://www.nature.com/articles/s41598-020-64665-2, doi: 10.1038/s41598-020-64665-2, publisher: Nature Publishing Group.

39. Kalidindi HT, Cross KP, Lillicrap TP, Omrani M, Falotico E, Sabes PN, Scott SH. Rotational dynamics in motor cortex are consistent with a feedback controller. eLife. 2021 Nov; 10:e67256. 10.7554/eLife.67256, doi: 10.7554/eLife.67256.

40. Kao TC, Sadabadi MS, Hennequin G. Optimal anticipatory control as a theory of motor preparation: A thalamocortical circuit model. Neuron. 2021 May; 109(9):1567–1581.e12. https://pmc.ncbi.nlm.nih.gov/articles/PMC8111422/, doi: 10.1016/j.neuron.2021.03.009.

41. Kaufman MT, Churchland MM, Ryu SI, Shenoy KV. Cortical activity in the null space: permitting preparation without movement. Nature Neuroscience. 2014 Mar; 17(3):440–448. doi: 10.1038/nn.3643.

42. Kaufman MT, Seely JS, Sussillo D, Ryu SI, Shenoy KV, Churchland MM. The Largest Response Component in the Motor Cortex Reflects Movement Timing but Not Movement Type. eNeuro. 2016 Jul; 3(4). https://www.eneuro.org/content/3/4/ENEURO.0085-16.2016, doi: 10.1523/ENEURO.0085-16.2016.

43. Kawai R, Markman T, Poddar R, Ko R, Fantana AL, Dhawale AK, Kampff AR, Ölveczky BP. Motor cortex is required for learning but not for executing a motor skill. Neuron. 2015 May; 86(3):800–812. doi: 10.1016/j.neuron.2015.03.024.

44. Kerlin A, Mohar B, Flickinger D, MacLennan BJ, Dean MB, Davis C, Spruston N, Svoboda K. Functional clustering of dendritic activity during decision-making. eLife. 2019 Oct; 8:e46966. https://elifesciences.org/articles/46966, doi: 10.7554/eLife.46966.

45. Krakauer JW, Hadjiosif AM, Xu J, Wong AL, Haith AM. Motor Learning. Comprehensive Physiology. 2019 Mar; 9(2):613–663. doi: 10.1002/cphy.c170043.

46. Kudryashova N, Hurwitz C, Perich MG, Hennig MH. BAND: Behavior-Aligned Neural Dynamics is all you need to capture motor corrections. bioRxiv. 2025 Mar; p. 2025.03.21.644350. https://pmc.ncbi.nlm.nih.gov/articles/PMC11974739/, doi: 10.1101/2025.03.21.644350.

47. Körding KP, König P. Supervised and Unsupervised Learning with Two Sites of Synaptic Integration. Journal of Computational Neuroscience. 2001 Nov; 11(3):207–215. 10.1023/A:1013776130161, doi: 10.1023/A:1013776130161.

48. Lacefield CO, Pnevmatikakis EA, Paninski L, Bruno RM. Reinforcement Learning Recruits Somata and Apical Dendrites across Layers of Primary Sensory Cortex. Cell Reports. 2019 Feb; 26(8):2000–2008.e2. https://linkinghub.elsevier.com/retrieve/pii/S2211124719301305, doi: 10.1016/j.celrep.2019.01.093.

49. Larkum ME, Kaiser KMM, Sakmann B. Calcium electrogenesis in distal apical dendrites of layer 5 pyramidal cells at a critical frequency of back-propagating action potentials. Proceedings of the National Academy of Sciences. 1999 Dec; 96(25):14600–14604. https://www.pnas.org/doi/abs/10.1073/pnas.96.25.14600, doi: 10.1073/pnas.96.25.14600.

50. Larkum M. A cellular mechanism for cortical associations: an organizing principle for the cerebral cortex. Trends in Neurosciences. 2013 Mar; 36(3):141–151. https://www.cell.com/trends/neurosciences/abstract/S0166-2236(12)00203-2, doi: 10.1016/j.tins.2012.11.006.

51. Larkum ME, Nevian T, Sandler M, Polsky A, Schiller J. Synaptic Integration in Tuft Dendrites of Layer 5 Pyramidal Neurons: A New Unifying Principle. Science. 2009 Aug; 325(5941):756–760. https://www.science.org/doi/full/10.1126/science.1171958, doi: 10.1126/science.1171958.

52. Larkum ME, Zhu JJ, Sakmann B. A new cellular mechanism for coupling inputs arriving at different cortical layers. Nature. 1999 Mar; 398(6725):338–341. https://www.nature.com/articles/18686, doi: 10.1038/18686.

53. Lecoq J, Oliver M, Siegle JH, Orlova N, Ledochowitsch P, Koch C. Removing independent noise in systems neuroscience data using DeepInterpolation. Nature Methods. 2021 Nov; 18(11):1401–1408. https://www.nature.com/articles/s41592-021-01285-2, doi: 10.1038/s41592-021-01285-2.

54. Lee BH, Park P, Wu X, Wong-Campos JD, Xu J, Xiong M, Qi Y, Huang YC, Itkis DG, Plutkis SE, Lavis LD, Cohen AE, Fast dendritic excitations primarily mediate back-propagation in CA1 pyramidal neurons during behavior. bioRxiv; 2026. https://www.biorxiv.org/content/10.64898/2026.01.03.696606v1, doi: 10.64898/2026.01.03.696606, iSSN: 2692-8205 Pages: 2026.01.03.696606 Section: New Results.

55. Levy S, Lavzin M, Benisty H, Ghanayim A, Dubin U, Achvat S, Brosh Z, Aeed F, Mensh BD, Schiller Y, Meir R, Barak O, Talmon R, Hantman AW, Schiller J. Cell-Type-Specific Outcome Representation in the Primary Motor Cortex. Neuron. 2020 Sep; 107(5):954–971.e9. https://linkinghub.elsevier.com/retrieve/pii/S0896627320304359, doi: 10.1016/j.neuron.2020.06.006.

56. Li N, Chen TW, Guo ZV, Gerfen CR, Svoboda K. A motor cortex circuit for motor planning and movement. Nature. 2015 Mar; 519(7541):51–56. https://www.nature.com/articles/nature14178, doi: 10.1038/nature14178.

57. Li N, Daie K, Svoboda K, Druckmann S. Robust neuronal dynamics in premotor cortex during motor planning. Nature. 2016 Apr; 532(7600):459–464. https://pmc.ncbi.nlm.nih.gov/articles/PMC5081260/, doi: 10.1038/nature17643.

58. Li Q, Ko H, Qian ZM, Yan LYC, Chan DCW, Arbuthnott G, Ke Y, Yung WH. Refinement of learned skilled movement representation in motor cortex deep output layer. Nature Communications. 2017 Jun; 8:15834. doi: 10.1038/ncomms15834.

59. Logiaco L, Abbott LF, Escola S. Thalamic control of cortical dynamics in a model of flexible motor sequencing. Cell Reports. 2021 Jun; 35(9):109090. doi: 10.1016/j.celrep.2021.109090.

60. Lopes G, Nogueira J, Dimitriadis G, Menendez JA, Paton JJ, Kampff AR. A robust role for motor cortex. Frontiers in Neuroscience. 2023 Feb; 17. https://www.frontiersin.org/journals/neuroscience/articles/10.3389/fnins.2023.971980/full, doi: 10.3389/fnins.2023.971980.

61. Makino H, Ren C, Liu H, Kim AN, Kondapaneni N, Liu X, Kuzum D, Komiyama T. Transformation of Cortex-wide Emergent Properties during Motor Learning. Neuron. 2017 May; 94(4):880–890.e8. https://www.sciencedirect.com/science/article/pii/S0896627317303410, doi: 10.1016/j.neuron.2017.04.015.

62. Malina KCK, Tsivourakis E, Kushinsky D, Apelblat D, Shtiglitz S, Zohar E, Sokoletsky M, Tasaka Gi, Mizrahi A, Lampl I, Spiegel I. NDNF interneurons in layer 1 gain-modulate whole cortical columns according to an animal’s behavioral state. Neuron. 2021 Jul; 109(13):2150–2164.e5. https://www.cell.com/neuron/abstract/S0896-6273(21)00327-5, doi: 10.1016/j.neuron.2021.05.001.

63. Malonis P, Vishnubhotla A, Hatsopoulos N, MacLean J, Kaufman M. Combatting nonidentifiability to infer motor cortex inputs yields similar encoding of initial and corrective movement. bioRxiv. 2021; 10.1101/2021.10.18.464704, doi: 10.1101/2021.10.18.464704.

64. Manita S, Suzuki T, Homma C, Matsumoto T, Odagawa M, Yamada K, Ota K, Matsubara C, Inutsuka A, Sato M, Ohkura M, Yamanaka A, Yanagawa Y, Nakai J, Hayashi Y, Larkum ME, Murayama M. A Top-Down Cortical Circuit for Accurate Sensory Perception. Neuron. 2015 Jun; 86(5):1304–1316. https://www.cell.com/neuron/abstract/S0896-6273(15)00413-4, doi: 10.1016/j.neuron.2015.05.006.

65. Mathis A, Mamidanna P, Cury KM, Abe T, Murthy VN, Mathis MW, Bethge M. DeepLabCut: markerless pose estimation of user-defined body parts with deep learning. Nature Neuroscience. 2018 Sep; 21(9):1281–1289. https://www.nature.com/articles/s41593-018-0209-y, doi: 10.1038/s41593-018-0209-y.

66. Mizes KGC, Lindsey J, Escola GS, Ölveczky BP. The role of motor cortex in motor sequence execution depends on demands for flexibility. Nature Neuroscience. 2024 Dec; 27(12):2466–2475. https://www.nature.com/articles/s41593-024-01792-3, doi: 10.1038/s41593-024-01792-3.

67. Moberg S, Takahashi N. Neocortical layer 5 subclasses: From cellular properties to roles in behavior. Frontiers in Synaptic Neuroscience. 2022 Oct; 14:1006773. https://www.frontiersin.org/articles/10.3389/fnsyn.2022.1006773/full, doi: 10.3389/fnsyn.2022.1006773.

68. Murray JD, Bernacchia A, Freedman DJ, Romo R, Wallis JD, Cai X, Padoa-Schioppa C, Pasternak T, Seo H, Lee D, Wang XJ. A hierarchy of intrinsic timescales across primate cortex. Nature Neuroscience. 2014 Dec; 17(12):1661–1663. https://www.nature.com/articles/nn.3862, doi: 10.1038/nn.3862.

69. Otor Y, Achvat S, Cermak N, Benisty H, Abboud M, Barak O, Schiller Y, Poleg-Polsky A, Schiller J. Dynamic compartmental computations in tuft dendrites of layer 5 neurons during motor behavior. Science. 2022 Apr; 376(6590):267–275. https://www.science.org/doi/10.1126/science.abn1421, doi: 10.1126/science.abn1421.

70. Pedregosa F, Varoquaux G, Gramfort A, Michel V, Thirion B, Grisel O, Blondel M, Prettenhofer P, Weiss R, Dubourg V, Vanderplas J, Passos A, Cournapeau D, Brucher M, Perrot M, Duchesnay E. Scikit-learn: Machine Learning in Python. J Mach Learn Res. 2011 Nov; 12(null):2825–2830. https://dl.acm.org/doi/10.5555/1953048.2078195.

71. Peng H, Xie P, Liu L, Kuang X, Wang Y, Qu L, Gong H, Jiang S, Li A, Ruan Z, Ding L, Yao Z, Chen C, Chen M, Daigle TL, Dalley R, Ding Z, Duan Y, Feiner A, He P, et al. Morphological diversity of single neurons in molecularly defined cell types. Nature. 2021 Oct; 598(7879):174–181. https://www.nature.com/articles/s41586-021-03941-1, doi: 10.1038/s41586-021-03941-1, publisher: Nature Publishing Group.

72. Pereira-Obilinovic U, Daie K, Chen S, Svoboda K, Darshan R. Neural dynamics outside task-coding dimensions drive decision trajectories through transient amplification. bioRxiv: The Preprint Server for Biology. 2025 Nov; p. 2025.11.20.689599. doi: 10.1101/2025.11.20.689599.

73. Perich MG, Conti S, Badi M, Bogaard A, Barra B, Rajan K, Bloch J, Courtine G, Micera S, Capogrosso M, Milekovic T. Motor cortical dynamics are shaped by multiple distinct subspaces during naturalistic behavior. bioRxiv. 2024 Jul; https://www.biorxiv.org/content/10.1101/2020.07.30.228767v3, doi: 10.1101/2020.07.30.228767.

74. Peters AJ, Lee J, Hedrick NG, O’Neil K, Komiyama T. Reorganization of corticospinal output during motor learning. Nature Neuroscience. 2017 Aug; 20(8):1133–1141. https://www.nature.com/articles/nn.4596, doi: 10.1038/nn.4596.

75. Petreanu L, Mao T, Sternson SM, Svoboda K. The subcellular organization of neocortical excitatory connections. Nature. 2009 Feb; 457(7233):1142–1145. https://www.nature.com/articles/nature07709, doi: 10.1038/nature07709.

76. Pnevmatikakis EA, Soudry D, Gao Y, Machado TA, Merel J, Pfau D, Reardon T, Mu Y, Lacefield C, Yang W, Ahrens M, Bruno R, Jessell TM, Peterka DS, Yuste R, Paninski L. Simultaneous Denoising, Deconvolution, and Demixing of Calcium Imaging Data. Neuron. 2016 Jan; 89(2):285–299. https://www.cell.com/neuron/abstract/S0896-6273(15)01084-3, doi: 10.1016/j.neuron.2015.11.037.

77. Pollak YE, Sachdev R, Larkum M, Gilad A. Cortex-wide laminar dynamics diverge during learning. bioRxiv. 2025 Jul; https://www.biorxiv.org/content/10.1101/2025.07.15.664840v1, doi: 10.1101/2025.07.15.664840.

78. Ranganathan GN, Apostolides PF, Harnett MT, Xu Nl, Druckmann S, Magee JC. Active dendritic integration and mixed neocortical network representations during an adaptive sensing behavior. Nature Neuroscience. 2018 Nov; 21(11):1583–1590. https://www.nature.com/articles/s41593-018-0254-6, doi: 10.1038/s41593-018-0254-6.

79. Rao RPN. A sensory–motor theory of the neocortex. Nature Neuroscience. 2024 Jul; 27(7):1221–1235. https://www.nature.com/articles/s41593-024-01673-9, doi: 10.1038/s41593-024-01673-9.

80. Riehle A, Requin J. Monkey primary motor and premotor cortex: single-cell activity related to prior information about direction and extent of an intended movement. Journal of Neurophysiology. 1989 Mar; 61(3):534–549. doi: 10.1152/jn.1989.61.3.534.

81. Rosier M, Stuyt G, Godenzini L, Ryan TJ, Palmer LM. Learning-induced plasticity decreases cortical engram cell dendritic excitability during memory recall. bioRxiv. 2025 Jan; https://www.biorxiv.org/content/10.1101/2025.01.08.631892v1, doi: 10.1101/2025.01.08.631892.

82. Sacramento J, Costa RP, Bengio Y, Senn W. Dendritic error backpropagation in deep cortical microcircuits. arXiv. 2017 Dec; http://arxiv.org/abs/1801.00062, doi: 10.48550/arXiv.1801.00062.

83. Sauerbrei BA, Guo JZ, Cohen JD, Mischiati M, Guo W, Kabra M, Verma N, Mensh B, Branson K, Hantman AW. Cortical pattern generation during dexterous movement is input-driven. Nature. 2020 Jan; 577(7790):386–391. https://www.nature.com/articles/s41586-019-1869-9, doi: 10.1038/s41586-019-1869-9.

84. Schoenfeld G, Kollmorgen S, Tsai MC, Lewis C, Han S, Bethge P, Reuss AM, Aguzzi A, Senn W, Mante V, Helmchen F. Unsigned temporal difference errors in cortical L5 dendrites during learning. bioRxiv. 2021 Dec; http://biorxiv.org/lookup/doi/10.1101/2021.12.28.474360, doi: 10.1101/2021.12.28.474360.

85. Scott SH. The computational and neural basis of voluntary motor control and planning. Trends in Cognitive Sciences. 2012 Nov; 16(11):541–549. https://linkinghub.elsevier.com/retrieve/pii/S1364661312002240, doi: 10.1016/j.tics.2012.09.008.

86. Senn W, Dold D, Kungl AF, Ellenberger B, Jordan J, Bengio Y, Sacramento J, Petrovici MA. A neuronal least-action principle for real-time learning in cortical circuits. eLife. 2024 Dec; 12:RP89674. 10.7554/eLife.89674, doi: 10.7554/eLife.89674.

87. Serradj N, Marino F, Moreno-López Y, Bernstein A, Agger S, Soliman M, Sloan A, Hollis E. Task-specific modulation of corticospinal neuron activity during motor learning in mice. Nature Communications. 2023 May; 14(1):2708. https://www.nature.com/articles/s41467-023-38418-4, doi: 10.1038/s41467-023-38418-4.

88. Shepherd GMG, Yamawaki N. Untangling the cortico-thalamo-cortical loop: cellular pieces of a knotty circuit puzzle. Nature Reviews Neuroscience. 2021 Jul; 22(7):389–406. https://www.nature.com/articles/s41583-021-00459-3, doi: 10.1038/s41583-021-00459-3.

89. Spratling MW. Cortical Region Interactions and the Functional Role of Apical Dendrites. Behavioral and Cognitive Neuroscience Reviews. 2002 Sep; 1(3):219–228. 10.1177/1534582302001003003, doi: 10.1177/1534582302001003003.

90. Stopfer M, Jayaraman V, Laurent G. Intensity versus Identity Coding in an Olfactory System. Neuron. 2003 Sep; 39(6):991–1004. https://www.cell.com/neuron/abstract/S0896-6273(03)00535-X, doi: 10.1016/j.neuron.2003.08.011.

91. Stuart GJ, Häusser M. Dendritic coincidence detection of EPSPs and action potentials. Nature Neuroscience. 2001 Jan; 4(1):63–71. https://www.nature.com/articles/nn0101_63, doi: 10.1038/82910.

92. Stuart GJ, Spruston N. Dendritic integration: 60 years of progress. Nature Neuroscience. 2015 Dec; 18(12):1713–1721. https://www.nature.com/articles/nn.4157, doi: 10.1038/nn.4157.

93. Takahashi N, Moberg S, Zolnik TA, Catanese J, Sachdev RNS, Larkum ME, Jaeger D. Thalamic input to motor cortex facilitates goal-directed action initiation. Current Biology. 2021 Sep; 31(18):4148–4155.e4. https://linkinghub.elsevier.com/retrieve/pii/S0960982221009180, doi: 10.1016/j.cub.2021.06.089.

94. Takahashi N, Oertner TG, Hegemann P, Larkum ME. Active cortical dendrites modulate perception. Science. 2016 Dec; 354(6319):1587–1590. doi: 10.1126/science.aah6066.

95. Todorov E, Jordan MI. Optimal feedback control as a theory of motor coordination. Nature Neuroscience. 2002 Nov; 5(11):1226–1235. https://www.nature.com/articles/nn963, doi: 10.1038/nn963.

96. Tran H, kerlin-lab/TStracker; 2026. https://github.com/kerlin-lab/TStracker.

97. Virtanen P, Gommers R, Oliphant TE, Haberland M, Reddy T, Cournapeau D, Burovski E, Peterson P, Weckesser W, Bright J, van der Walt SJ, Brett M, Wilson J, Millman KJ, Mayorov N, Nelson ARJ, Jones E, Kern R, Larson E, Carey CJ, et al. SciPy 1.0: fundamental algorithms for scientific computing in Python. Nature Methods. 2020 Mar; 17(3):261–272. https://www.nature.com/articles/s41592-019-0686-2, doi: 10.1038/s41592-019-0686-2.

98. Walt Svd, Schönberger JL, Nunez-Iglesias J, Boulogne F, Warner JD, Yager N, Gouillart E, Yu T. scikit-image: image processing in Python. PeerJ. 2014 Jun; 2:e453. https://peerj.com/articles/453, doi: 10.7717/peerj.453.

99. Wang Q, Ding SL, Li Y, Royall J, Feng D, Lesnar P, Graddis N, Naeemi M, Facer B, Ho A, Dolbeare T, Blanchard B, Dee N, Wakeman W, Hirokawa KE, Szafer A, Sunkin SM, Oh SW, Bernard A, Phillips JW, et al. The Allen Mouse Brain Common Coordinate Framework: A 3D Reference Atlas. Cell. 2020 May; 181(4):936–953.e20. https://www.sciencedirect.com/science/article/pii/S0092867420304025, doi: 10.1016/j.cell.2020.04.007.

100. Wu X, Lee BH, Park P, Wong-Campos JD, Xu J, Plutkis SE, Lavis LD, Cohen AE, A dendrite-resolved, in vivo transfer function from spike patterns to dendritic Ca2+. bioRxiv; 2026. https://www.biorxiv.org/content/10.64898/2026.01.18.700189v1, doi: 10.64898/2026.01.18.700189, iSSN: 2692-8205 Pages: 2026.01.18.700189 Section: New Results.

101. Xiao K, Li Y, Sullivan BJ, Li G, Magee JC. Rapid neocortical network modifications via dendritic plateau potential induced plasticity. bioRxiv. 2025 Nov; https://www.biorxiv.org/content/10.1101/2025.11.19.689338v1, doi: 10.1101/2025.11.19.689338.

102. Xu Nl, Harnett MT, Williams SR, Huber D, O’Connor DH, Svoboda K, Magee JC. Nonlinear dendritic integration of sensory and motor input during an active sensing task. Nature. 2012 Dec; 492(7428):247–251. https://www.nature.com/articles/nature11601, doi: 10.1038/nature11601.

103. Xu T, Yu X, Perlik AJ, Tobin WF, Zweig JA, Tennant K, Jones T, Zuo Y. Rapid formation and selective stabilization of synapses for enduring motor memories. Nature. 2009 Dec; 462(7275):915–919. https://www.nature.com/articles/nature08389, doi: 10.1038/nature08389.

104. Yaeger CE, Mojica Soto-Albors R, Liu W, Beltramini A, Harnett MT. Plateau potentials are instructive signals for behavioral timescale synaptic plasticity in the neocortex. bioRxiv: The Preprint Server for Biology. 2025 Nov; p. 2025.11.07.687250. doi: 10.1101/2025.11.07.687250.

105. Yang W, Tipparaju SL, Chen G, Li N. Thalamus-driven functional populations in frontal cortex support decision-making. Nature Neuroscience. 2022 Oct; 25(10):1339–1352. https://www.nature.com/articles/s41593-022-01171-w, doi: 10.1038/s41593-022-01171-w.

106. Zhang Y, Rózsa M, Liang Y, Bushey D, Wei Z, Zheng J, Reep D, Broussard GJ, Tsang A, Tsegaye G, Narayan S, Obara CJ, Lim JX, Patel R, Zhang R, Ahrens MB, Turner GC, Wang SSH, Korff WL, Schreiter ER, et al. Fast and sensitive GCaMP calcium indicators for imaging neural populations. Nature. 2023 Mar; 615(7954):884–891. https://www.nature.com/articles/s41586-023-05828-9, doi: 10.1038/s41586-023-05828-9.

107. Zhang Y, Liu Y, Hu Y, Zhang Z, Du J, Cui L, Liu Q, Zhong L, Xin Y, Chai R, Deng L, Pan J, Schiller J, Xu Nl. Dendritic computation for rule-based flexible categorization. bioRxiv. 2025 Jun; doi: 10.1101/2025.06.03.657766.

